# Spatiotemporal Patterning enabled by Gene Regulatory Networks

**DOI:** 10.1101/2022.04.13.488152

**Authors:** Ushasi Roy, Divyoj Singh, Navin Vincent, Chinmay Haritas, Mohit Kumar Jolly

## Abstract

Spatiotemporal pattern formation plays a key role in various biological phenomena including embryogenesis and neural network formation. Though the reaction-diffusion systems enabling pattern formation have been studied phenomenonlogically, the biomolecular mechanisms behind these processes has not been modelled in detail. Here, we study the emergence of spatiotemporal patterns due to simple synthetic commonly observed two- and three-node gene regulatory network motifs coupled with their molecular diffusion in one- and two-dimensional space. We investigate the patterns formed due to the coupling of inherent multistable and oscillatory behavior of toggle switch (two mutually repressing nodes), toggle switch with double self-activation, toggle triad (three mutually repressing nodes) and repressilator (three nodes repressing the other sequentially in a cyclic manner) with the effect of spatial diffusion of these molecules. We probe various parameter regimes corresponding to different regions of stability (monostable, multistable, oscillatory) and assess the impact of varying diffusion coefficients too. This analysis offers valuable insights into the design principles of pattern formation facilitated by these network motifs, and suggest mechanistic underpinnings of biological pattern formation.

## 1 Introduction

Pattern formation in living organisms is ubiquitous in nature, for instance, during embryonic development [1], morphogenesis [2], organization of neural networks [3, 4] and patterns on body (wing/skin/fur) [5]. In smaller organisms, we observe emergence of patterns during formation of various structures, in Hydra [6] and in the different species of *Drosophila* [1]. Some examples in higher organisms include wing decoration in butterfly, stripes in zebras, patches in giraffes or spots in leopards. These diverse patterns are often driven by morphogen gradients, and have been extensively investigated [7–9]. However, the underlying mechanisms in terms of gene regulatory networks are not as well-investigated. Thus, reproducing these patterns in synthetic biology has remained a challenging problem.

The theory of pattern formation has been explored mathematically since 1952 by Alan Turing in his seminal work [10]. In 1958, spontaneous, dynamic and oscillating concentric circular and spiral chemical patterns were first observed experimentally, famously known as, Belousov-Zhabotinsky (BZ) reactions [11, 12]. In 1972, Gierer and Meinhardt [13] developed a theoretical model of the Activator-Inhibitor system, based on Turing’s theory of 1952 [10], and experimentally designed it later [14–16]. In this two-component system, designed experimentally, an activator triggers the inhibitor while the inhibitor represses the activator in a long-range way. This mechanism could model different patterns including parallel stripes in shells, caused by near synchronous oscillations. Other theoretical models [17–19] generated similar patterns. Thus, Turing’s work in 1952 played a key role in driving theoretical work applied to biological systems [10].

Mathematical models incorporating diffusion of one of the two components have been developed to understand the underlying mechanisms of various systems [20, 21]. Moreover, experiments have been designed in various contexts of single diffusing molecule [22, 23], spatial arrangement and manipulation of inducers [24, 25], and quorum sensing molecules [26, 27], again highlighting the importance of Turing model in exploring pattern formation in many systems [17–19]. However, patterns emerging from diffusing molecules of gene regulatory networks is still an area of study in its formative stages and there are many challenges to it [28].

The inherent stochasticity due to spatial diffusion of molecules enhances the noise/fluctuations in the environment of a cell which can influence gene expression [29]. Molecular diffusion plays an important role in transcriptional regulation, precision in biochemical signaling etc. [30]. Cottrell *et al*. developed an analytical model for branching-diffusion and estimated the mean and the magnitude of fluctuations which can be determined by the protein’s Kuramoto length — the typical distance across which a protein can diffuse in its lifetime [31]. How such diffusion coupled with emergent dynamics of the gene regulatory networks can generate specific patterns, remains relatively less explored [28, 32–34].

Here, we aim to study various diverse patterns emerging out of spatiotemporal dynamics of interactive diffusive molecules of different simple two- and three-node gene regulatory network (GRN) motifs – toggle switch [35] with/without self-activation [36], toggle triad [37] and repressilator [38]. We have further observed how the patterns change corresponding to different stability regimes of a single motif, and for varying diffusion coefficients. The patterns induced by GRNs are transient, unstable, evolve dynamically, and asymptotically reach a stable homogeneous state. The dynamics of pattern formation depend on the motif being investigated and operating regimes (monostable, oscillatory, multistable).

## 2 Results

### 2.1 Toggle Switch - an archetype of GRNs - exhibiting temporal and spatial bistability

We take a bottom-up approach and begin our study by analyzing a toggle switch, a network formed by two genes *A* and *B*, mutually repressing each other (schematic shown in the inset of the first plot of Fig.1 A). Each of *A* and *B* is usually a master regulator of a specific cellular fate and inhibits the activity of the other gene. It is an archetype of GRNs in various biological systems, for instance, mammalian cell fate tree (with toggle switch at each of its branch point) [39].A toggle switch enables two states: (high A, low B) which we call as state A and (low A and high B) which we call as state B. It displays both monostable and bistable behaviors at different parameter regimes. We can find bistable behaviors in a variety of biological systems, such as embryonic patterning of Drosophila [2, 40] or signaling networks [41] or galactose regulatory networks [42, 43].

**Figure 1.**
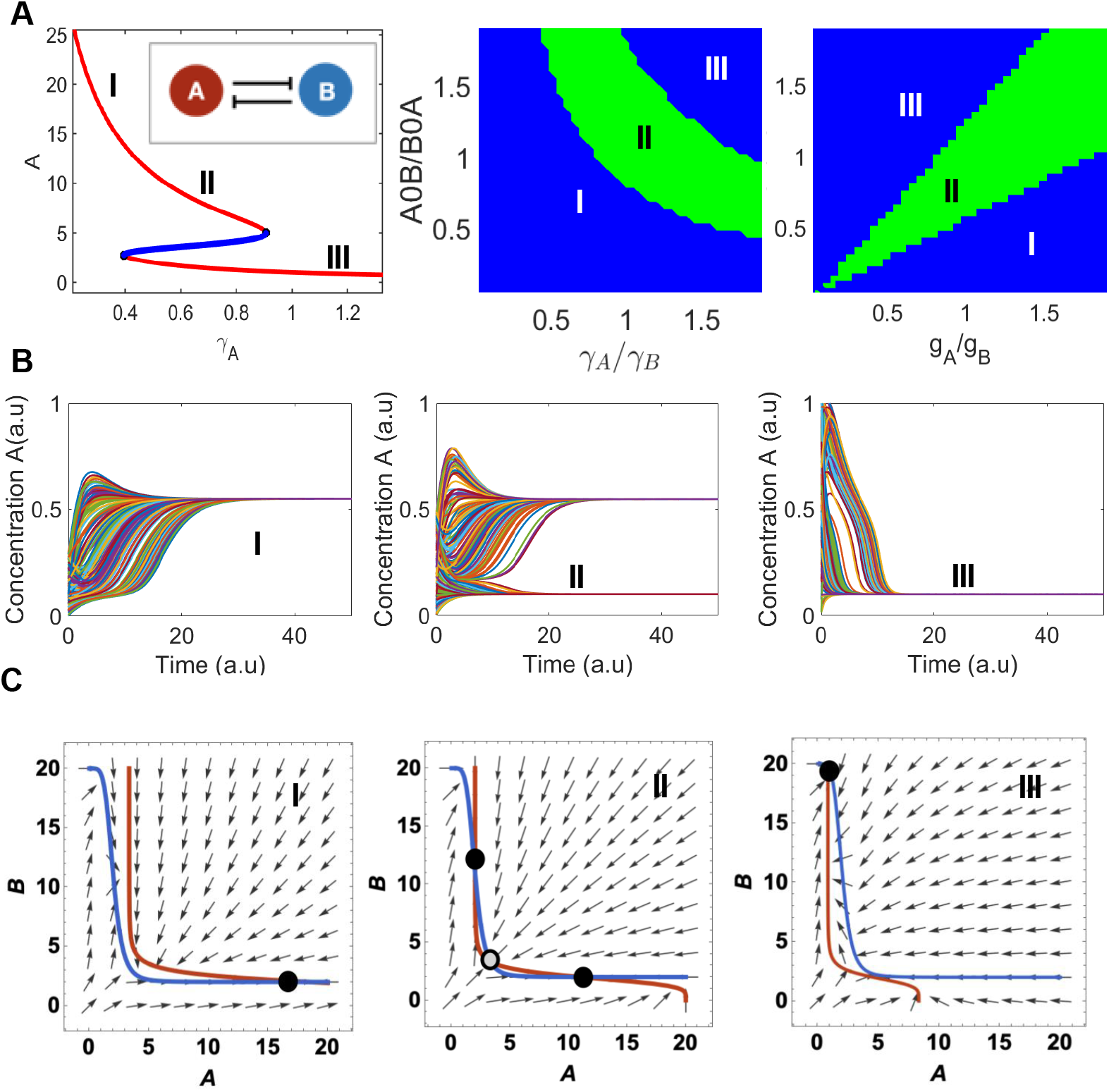
Temporal dynamics of Toggle Switch (TS) A) Left - Bifurcation diagram of one of the proteins (A) as a function of degradation rate (*γ*_*A*_) depicting the bistable behavior of a toggle switch (inset); Centre and - Right Phase Diagrams between non-dimensionliased parameters which control the strength of toggle switch. Different colors depict different regions of bistability and monostability. *A*0*B/B*0*A, γ*_*A*_*/γ*_*B*_, *g*_*A*_*/g*_*B*_ are the ratio of thresholds, degradation rates and production rate, respectively. B) Time evolution of one of the proteins (A here) starting from different initial conditions under different regimes of bifurcation diagram C) Nullclines for different regimes of the parameter space.

The governing equation for genetic toggle switch - two genes mutually repressing each other - in terms of the shifted Hill function ℋ is given by

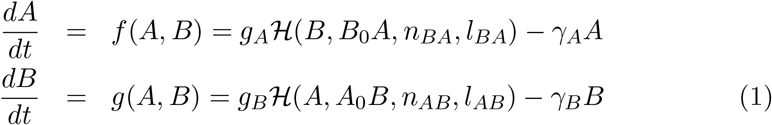

where *g*_*A*_ and *g*_*B*_ are the production rates, *γ*_*A*_ and *γ*_*B*_ are the degradation rates. The first terms in the equations (1) correspond to the effective production rates of the species and the second terms corresponds to the degradation of the species. The terms ℋ (*B, B*_0_*A, n*_*BA*_, *l*_*BA*_) and ℋ (*A, A*_0_*B, n*_*AB*_, *l*_*AB*_) are the shifted Hill functions and capture the interaction between gene A and gene B due to transcriptional inhibition.

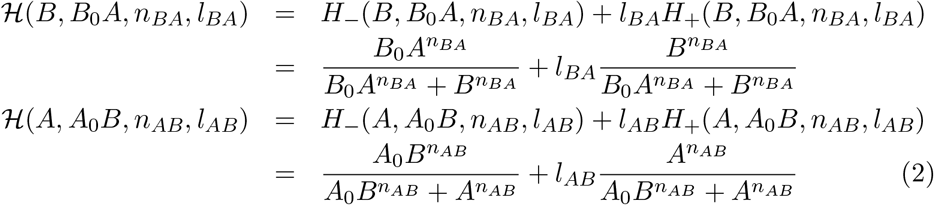

In (2) *B*_0_*A* and *A*_0_*B* are the thresholds which determine the levels at which the transcriptional interaction becomes active. The Hill coefficients, *n*_*BA*_ and *n*_*AB*_ correspond to the cooperativity of interaction and the *l*_*BA*_ and *l*_*AB*_ are the fold change due in the levels of proteins due to transcriptional activation or inhibition. The fold change value of *l*_*BA*_ *>* 1 means that B is activating A and *l*_*BA*_ < 1 means that B is inhibiting A [44].

First, we perform numerical analysis to understand the dynamical behavior of the toggle switch motif. Numerical bifurcation analysis shows that the system switches from a monostable region of state A (high A, low B Ab) to a bistable region where both state A (high A, low B Ab) and state B (low A, high B) are possible, to another monostable region with state B (low A, high B) (Fig. 1 A. The bifurcation parameter here is the degradation rate of A (*γA*). Starting from state A (region I), as we increase the degradation rate, the steady state concentration of A decreases and the system reaches a bistable region (region II), where the system can achieve both the states - A or B. If we further increase the degradation rate of A, the system can only attain state B (where B is high and A is low). Opposite behaviour can be seen if the degradation rate of B is chosen as a bifurcation parameter (Fig. S1 B).

Next, we draw the phase diagram depicting the stability regions. On the y-axis, we have *A*0*B/B*0*A* and as we increase this ratio, the relative strength of inhibition of A on B as compared to B on A, decreases. In the first phase diagram, we have the ratio of their degradation rates (*γA/γB*) on x-axis. As this ratio increases, the inhibition strength by A decreases. Therefore, the region marked I is the region where we observe state A and in the region II, the inhibition of both A and B are balanced and we observe a bistable behaviour. In the region III, the inhibition of A on B is weak as compared to B’S inhibition on A, therefore we observe the state B (high B, low A Ba)

Next, we numerically solved the equations (1) starting from 200 different initial conditions and from parameters in different regions. Dynamics of corresponding region reveal the system converging to one or both states (Fig. 1 B). The dynamics of B are presented in Fig. S1 A-B.

Panel C depicts the nullclines for the three regimes having one, two and one stable fixed points marked by solid black circles. The unstable steady state is shown by gray circle. The red curve is the plot for the solution of the first one of Eq.1 in the steady state while the blue curve is the same for the second one of Eq.1. The arrows represent the vector fields. They are directed towards the stable fixed point and away from the unstable one. A monostable regime is signified by having a single stable steady state or cells having one phenotype while a bistable regime signifies the possibility of a population of cells having two different fates denoted by two stable steady states. The bistable temporal dynamics of gene B for parameter set 1 corresponding to three stability regimes (same as in Fig. 1) and its bifurcation diagram, the phase diagrams (ratio of thresholds as a function of ratios of production rates and degradation rates respectively) and nullclines for parameter sets 2 and 3 are shown in Fig. S1 D. Phase diagrams of the time taken for gene A to reach the steady state are also presented in the Fig. S2 A-C.

#### 2.1.1 Toggle Switch with 1D diffusion

Tkacik and Bialek developed a mathematical model for understanding the one-dimensional diffusion - longitudinal sliding of repressor molecules along DNA during transcriptional regulation [30]. Here, we consider a one-dimensional chain of cells (depicting hair-line/filamentous structures), each having a toggle switch with diffusing gene molecules (see Fig.2A for the schematic). Each molecule can diffuse through the entire one-dimensional space. The mathematical equations describing the system can be written in the following way,

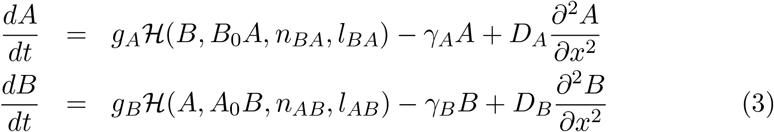

**Figure 2.**
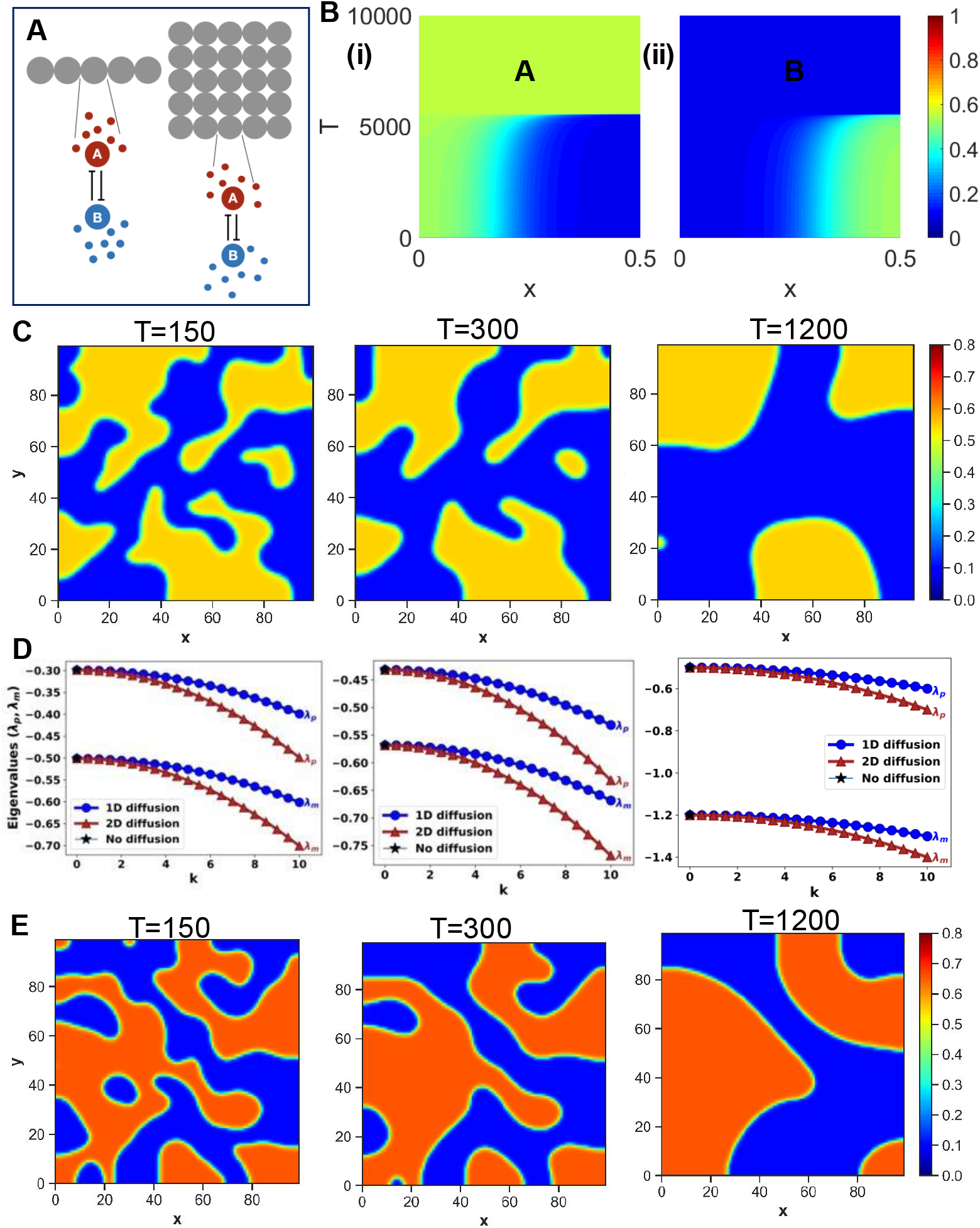
Spatiotemporal dynamics of Toggle Switch. A) Schematic of toggle switch plus diffusion system. In the 1D system, we consider a single array of cells and each of them have a toggle switch between diffusive species. In the 2D system, we have a square lattice of cells with diffusive species. B) Heatmaps of levels of protein A (i) and B (ii) as a function of time and space in the 1D diffusive system. C) Heatmap of levels of A in the 2D diffusive system at different time points D) Analytical plots in different parameter regimes, viz. Monostable-I, Bistable-II and Monostable-III (refer to previous figure). E) Heatmaps of levels of A for another parameter set (Parameter set-2) at different time points.

The above equations are similar to 1 with one added term in the right hand side describing diffusion. *D*_*A*_ and *D*_*B*_ are the diffusion constants/diffusivities for gene A and B. Higher diffusivity refers to faster diffusion. Fig.2 B shows bistable spatiotemporal behavior of toggle switches with molecules diffusing in one dimensional space. Green and blue represent the higher and the lower expression levels of gene A. The simulation starts with a spatially in-homogeneous state (having both expression levels) but as time progresses, the system reaches its steady homogeneous state. One can notice the complimentary behavior of gene (i) *A* and (ii) *B* as expected from a toggle switch.

#### 2.1.2 Toggle Switch with 2D diffusion

Consider a two-dimensional square lattice of cells (depicting thin mono-layer of biological tissue), each having a toggle switch with diffusing gene molecules (see Fig.2A for the schematic). Each molecule can diffuse through the entire two-dimensional space.

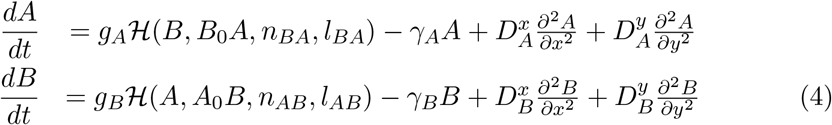

In two dimensions, the equations have two added terms in the right hand side describing diffusion in two different spatial direction. 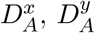 and 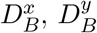 are the diffusion constants/diffusivities for gene A and B in the x- and y-direction respectively.

The spatiotemporal behavior of toggle switches with molecules diffusing in two dimensional space at three different time instants is shown in Fig. 2 C. We notice that as time progresses, the size of the patches increases (Suppl Videos S3.1). Panel D shows analytical plots for eigenvalues in different parameter regimes, viz. Monostable-I, Bistable-II and Monostable-III. Analytical solutions reveal negative eigenvalues, thus indicating the steady-ness of the patterns. With the increase in wavenumber *k*, eigenvalues become more negative. For *k* = 0, the system reduces to diffusion-free cases. In two dimensions, the fast convergence to underlying patterns, as seen across multiple parameter sets, further strengthen the stability and robustness of observed patterns. We also conducted similar analysis for another parameter set (Fig. 2 E) and found that the patterns are similar. The stability (quantified by the nature of trace, determinant and eigenvalues), the patterns in one-dimension is similar and robust across parameters (shown for parameter set 1, 2 and 3 in Fig. S3 A-C and S4 A-C).

### 2.2 Toggle Switch with self-activation - exhibiting spatiotemporal tristability

The next motif we consider is two self-activation loops coupled with a toggle switch (Fig.3 A). This architecture introduces a third stable steady state in the system. The balance between mutual repression and double self-activation enhances the probability of having a third state (intermediate A - intermediate B) in addition to the two possibilities of expression levels of toggle switch (high A - low B) and (low A - high B) [36, 37, 45, 46].

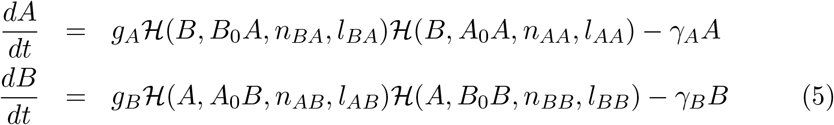

where ℋ(*B, B*_0_*A, n*_*BA*_, *l*_*BA*_) and ℋ(*A, A*_0_*B, n*_*AB*_, *l*_*AB*_) are the shifted Hill functions as given in Eq.(2) in the previous section, while the other two shifted Hill functions are for the self loops, given by the following,

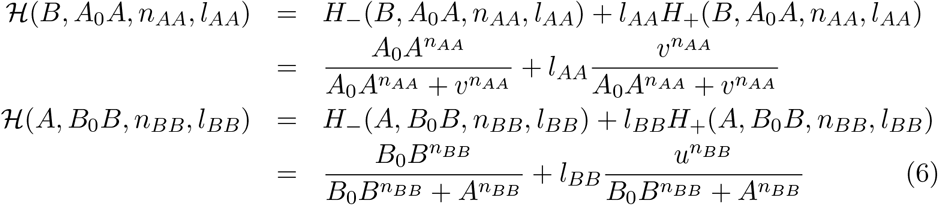

**Figure 3.**
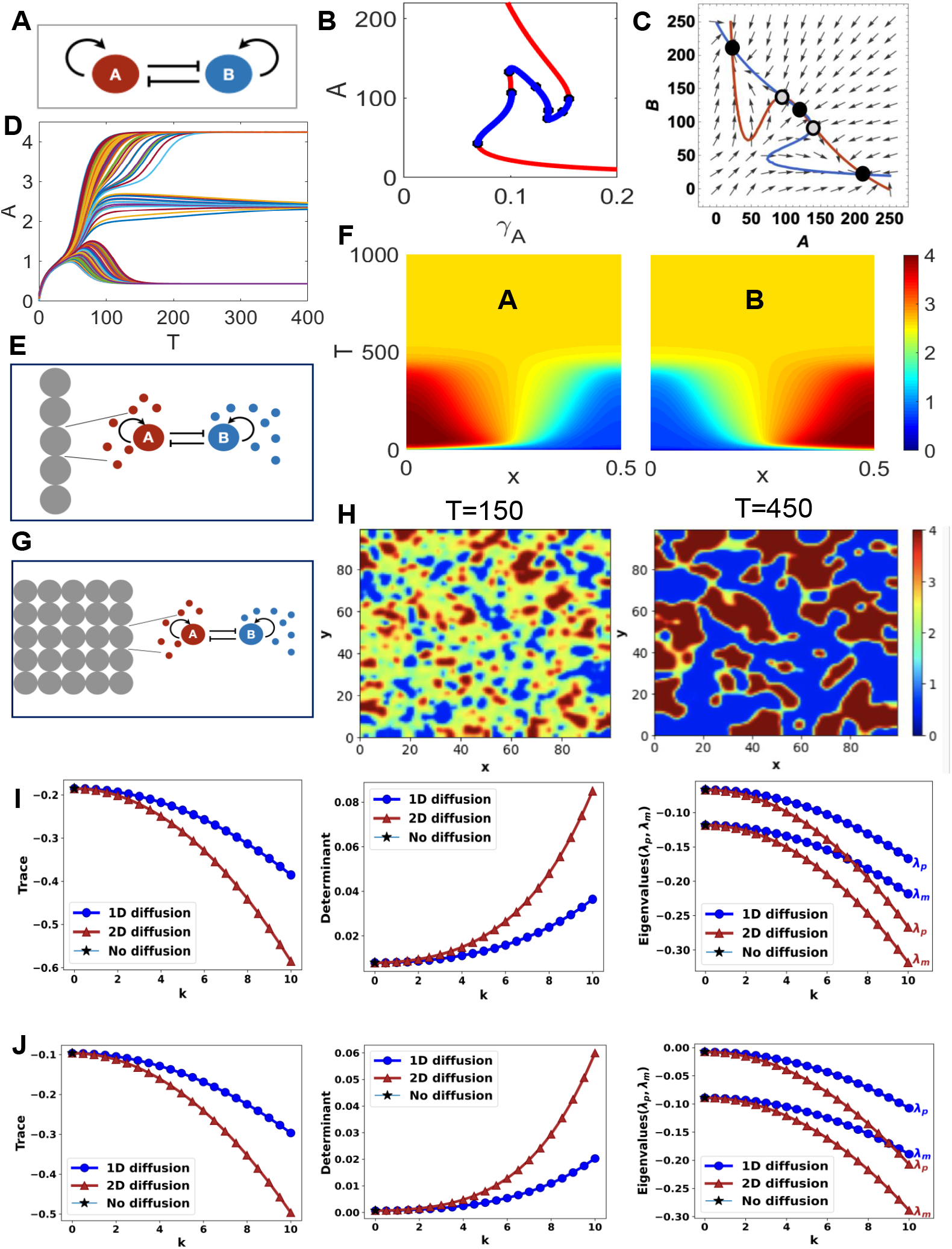
Dynamics of Toggle Switch with Self Activation (TSSA): A) Schematic of the ODE model of toggle switch between two species in a cell and self activation on both B) Bifurcation diagram of A as a function of degradation rate of A. Note the presence of a tristable region. C) Nullcline plot of the tristable region. D) Time evolution plot of protein A for multiple initial conditions in the tristable parameter regime. E) Schematics of the TSSA coupled with the 1D diffusive system. F) Heatmap of levels of A and B as a function of space and time for parameter set A G) Schematics of the TSSA coupled with the 2D diffusive system H) Heatmaps of levels of A for parameter set 1 at two different time points. I) Trace, determinants and eigenvalues evaluated at equilibrium points 1 and 3 J) Trace, determinants and eigenvalues evaluated at equilibrium points 2

Fig.3 B shows the bifurcation diagram of A. The expression levels of A is shown as a function of degradation rate of A, *γ*_*A*_. Three stable states are shown in red, while two unstable states are shown in blue. The corresponding nullclines are shown in (Fig.3 C). The three stable fixed points are marked by solid black circles while the two unstable steady states are shown by gray circles. (Fig.3 D) shows the time evolution of gene A for 200 different initial conditions, finally all ending up in three different stable steady states.

#### 2.2.1 Toggle Switch with self-activation with 1D diffusion

Let us now consider a one-dimensional chain of cells (depicting hair-line/filamentous structures), each having a toggle switch coupled with two self-activation loops (Schematic in Fig. 3 E). Each molecule can diffuse through the entire one-dimensional space. The governing equations are

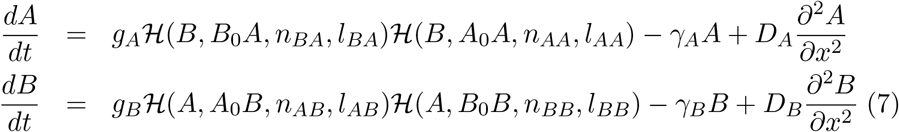

where ℋ are the shifted Hill functions as described in the previous sections and the last two terms in the RHS describe molecular diffusion (*D*_*A*_ and *D*_*B*_ being the diffusion coefficient of A and B respectively). Fig. 3 F shows the heat map of expression levels of A and B as a function of space and time for parameter set 1. The complimentarity in expression levels of gene A and gene B are shown side by side for comparison.

#### 2.2.2 Toggle Switch with self-activation with 2D diffusion

Consider a two-dimensional square lattice of cells (depicting thin mono-layer of biological tissue), each having a toggle switch coupled with two self-activation loops (Fig. 3 G). Each molecule can diffuse through the entire two-dimensional space. The mathematical equations describing the scenario are given by

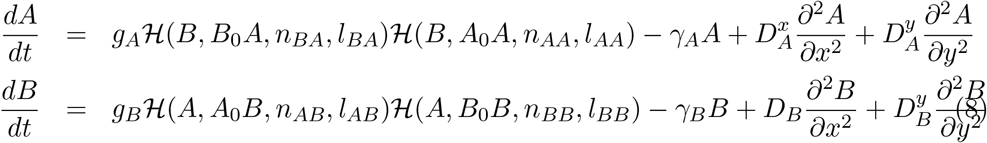

where the shifted Hill functions are expressed same as in the previous section, and there are two diffusive terms (in x- and y- spatial directions) for each gene.

Fig. 3 H shows the heatmaps of levels of gene A for parameter set 1 at two different time points for TSSA with molecules diffusing in two-dimensional space. We observe three states in the initial time point (T=150) A high, intermediate or low. As time progresses and molecules diffuse more, the intermediate state vanishes A-high or A low are observed dominantly (Suppl Video S3.2). In Fig. 3 I, we show the analytical plots for the trace, determinants, and eigenvalues evaluated at equilibrium points 1 (we obtain exactly same results are obtained at the second equilibrium point since the coordinates of equilibrium points are in symmetric pairs) and J shows the same quantities evaluated at equilibrium points 2. The temporal evolution and the spatiotemporal dynamics corresponding to tristable parameter regime is shown here. We also solved the system in a bistable parameter regime studied the steadys-state and spatio-temporal dynamics in supplementary (Fig. S5 A -G). The overall behaviour is very similar to bistable regime in toggle switch.

### 2.3 Toggle Triad - a frustrated tristable system

We consider a toggle triad, three coupled toggle switches, comprising the genes *A, B*, and *C* mutually repressing each other (schematic shown in Figure 4 A). This motif can have ‘pure’/single positive (A-high, B-low, C-low {Abc}) or (A-low, B-high, C-low {aBc}) or (A-low, B-low, C-high {abC}), ‘hybrid’/double positive (A-high, B-high, C-low {ABc}) or (A-low, B-high, C-high {aBC}) or (A-high, B-low, C-high {AbC}) or triple positive (A-high, B-high, C-high {ABC}) and triple negative (A-low, B-low, C-low {abc}) states.In our previous study [37], we showed that the frequency of triple positive and triple negative states is very low. Single positive states are more frequent compared to the double positive states. It should be noted that toggle triad is a frustrated system [47]; e.g. high levels of A will drive levels B to be low by repressing it, and low levels of B will make levels of C higher, that will eventually reduce the levels of A.

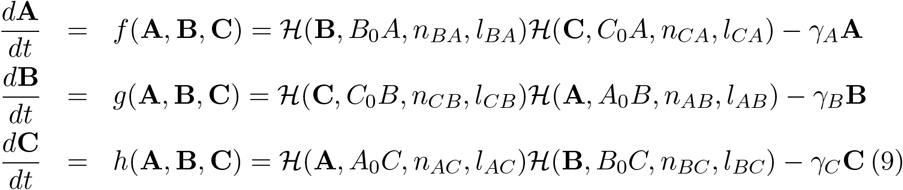

where ℋ are the shifted Hill functions defined as in Equations with different arguments which can be obtained by systematic changes. Here we focus on monostability, bistability, and tristability of single positive states since they are more frequent. The patterns formed by double-positive states are also not significantly different from the single-positive ones.

**Figure 4.**
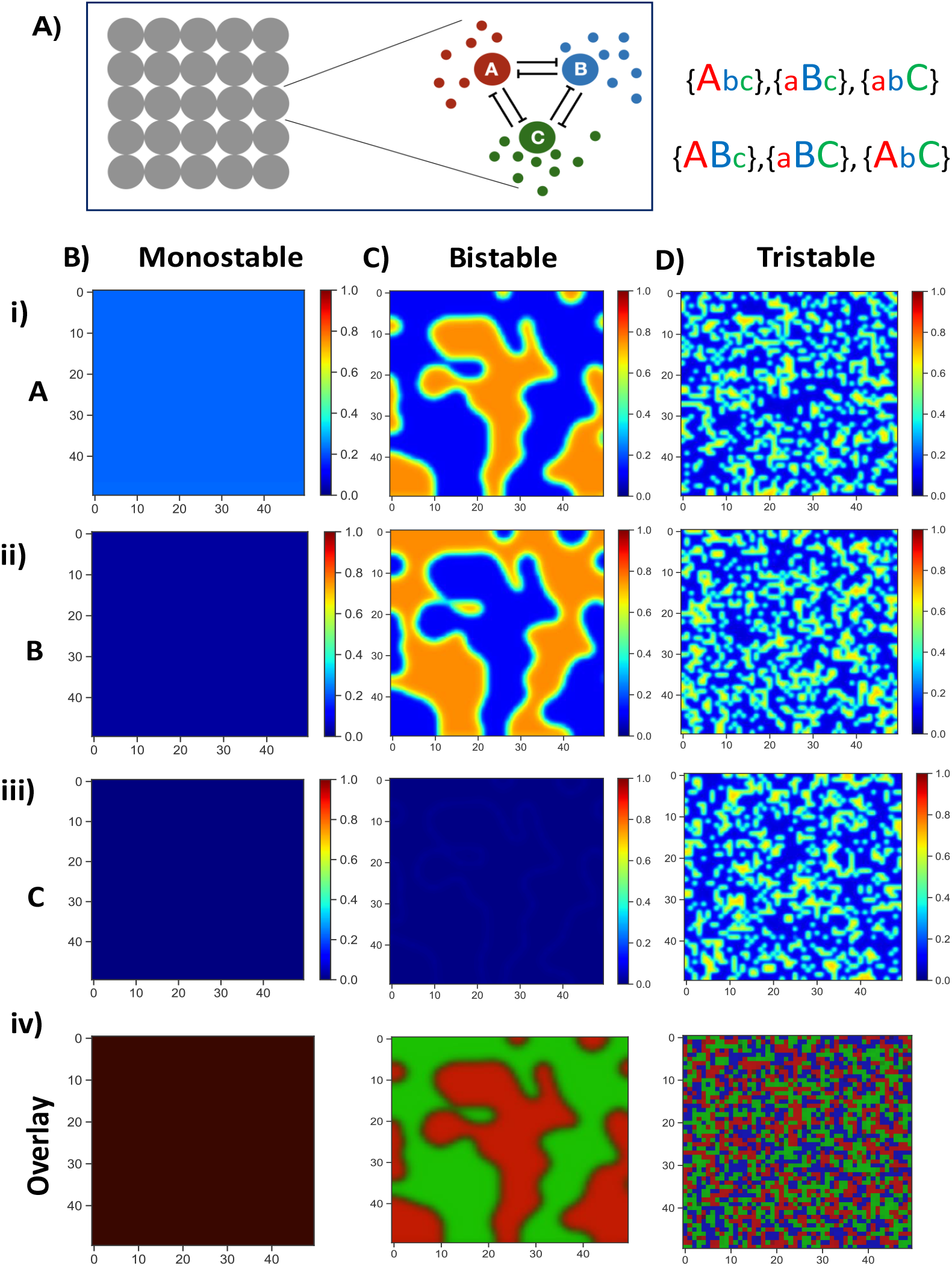
Dynamics of Toggle Triad. A) Schematic of toggle triad of three species in a cell coupled with the diffusion system. B) Heat map of levels of i) A ii) B iii) C as a function of space in the monostable parameter regime. Protein A is high in all cells as compared to B and C. iv) RGB (Red Green Blue) plot of the 3 species combined. Here, red corresponds to A high, green corresponds to B high, and blue corresponds to C high. C) Same as B) for the bistable parameter regime where either A is high or B is high. D) Same as B) but for tristable parameter regime in which either A is high or B is high or C is high.

#### 2.3.1 Toggle Triad with 2D diffusion

To study the pattern formation due to toggle triad, we coupled the motifs into the 2D lattice of cells. We write reaction-diffusion equations as follows:

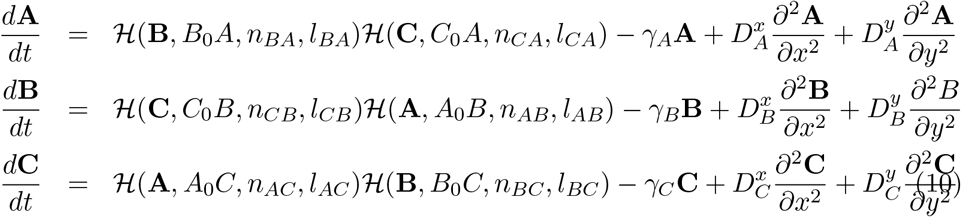

We first explore the evolving patterns in different phases, i.e, monostable, bistable, and tristable. When the motif shows the monostable behaviour (here A-high, B-low and C-low {Abc}), the system reaches a homogeneous state (Figure 4 B), as expected. When the motif is in the bistable parameter regime (here {Abc}-{aBc}), we get domains of regions where A is high and others where B is high. C remains low in all regions (Figure 4 C). In the tristable parameter regime (here {Abc}-{aBc{abC}), we pattern formation in all three species. The 2D space is distributed into different regions where either A is high or B is high or C is high. From the overlay plot is clear that only one of the genes is high in each cell (Figure 4 D) reflecting patterns seen in toggle triad (Suppl Videos S3.3).

Next, we studied the role of diffusion constant on the pattern formation. In the tristable parameter regime, when the diffusion constant is low (*D* = 0.0008), we observe that the patch size of regions where either of the three state (Abc, aBc or abC) is small (Fig. 5 A) as compared to the case where diffusion constant is intermediate (*D* = 0.008) (Fig. 5 B which is same as Fig. 4 D). As the diffusion constant increases (*D* = 0.08), the patch size of the patterns increases(Fig. 5C).

**Figure 5.**
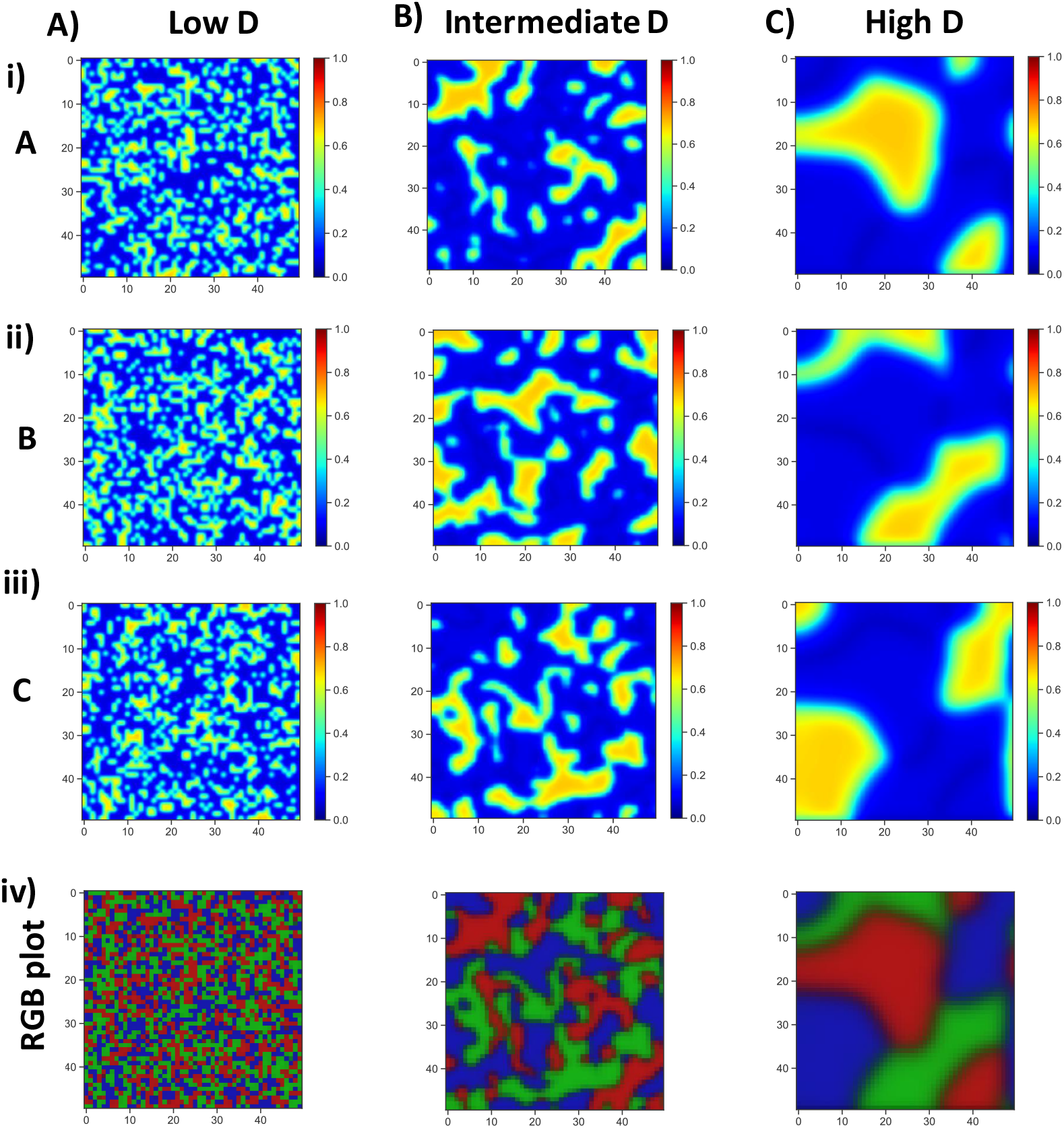
Dynamics of Toggle Triad in the tristable parameter regime for different diffusion constants. A) Heat map of levels of i) A ii) B iii) C as a function of space in the tristable parameter regime when the diffusion is low (D=0.0008). iv) RGB plot of the 3 species combined. Here, red corresponds to A high, green corresponds to B high, and blue corresponds to C high. B) Same as A but for intermediate diffusion levels (D=0.008) C) Same as A but for high diffusion levels (D=0.08)

Finally, we also studied the effect of variable diffusion on the pattern formation from a toggle triad motif in bistable regime (see Figure S6 A-C of SI).

Time evolution (Panel A and C) of the molecules of gene A for multiple initial conditions and the frequency distributions (Panel B and D) of different states at steady state in the bistable and tristable parameter regimes of toggle triad is shown in Fig. S7. Similarly as in the cases for previous motifs, the stabilities of the patterns are quantified by traces, determinants and three eigenvalues for different stability (mono (Fig. S8), bi (Fig. S9) and tri (Fig. S10)) regimes and for low (Panel A) and high (Panel B) diffusivity.

### 2.4 Repressilator - showcasing spatiotemporal oscillations

Oscillatory/rhythmic behaviors in biological systems are often generated by the complex interactions among various genes, proteins, metabolites etc. A number of factors like negative feedback, time delay, inherent nonlinearity of the reaction kinetics can give rise to biochemical oscillations [48]. Elowitz and Leibler [38] first designed and constructed an artificial biological clock, called repressilator, with three transcription factors in *E. coli*.

By weakening three links of Toggle Triad (see schematic in (Figure 6 A) and corresponding parameter set in the excel sheet), we obtain the motif for repressilator genes, *A, B* and *C* repress each other in a cyclic fashion. This is a well-known example of a biological oscillator. The governing equations are given by

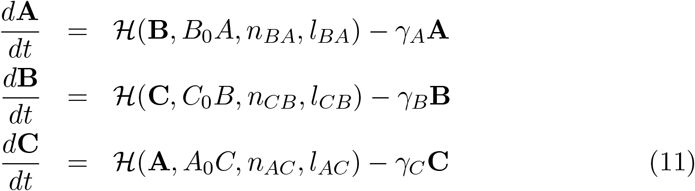

**Figure 6.**
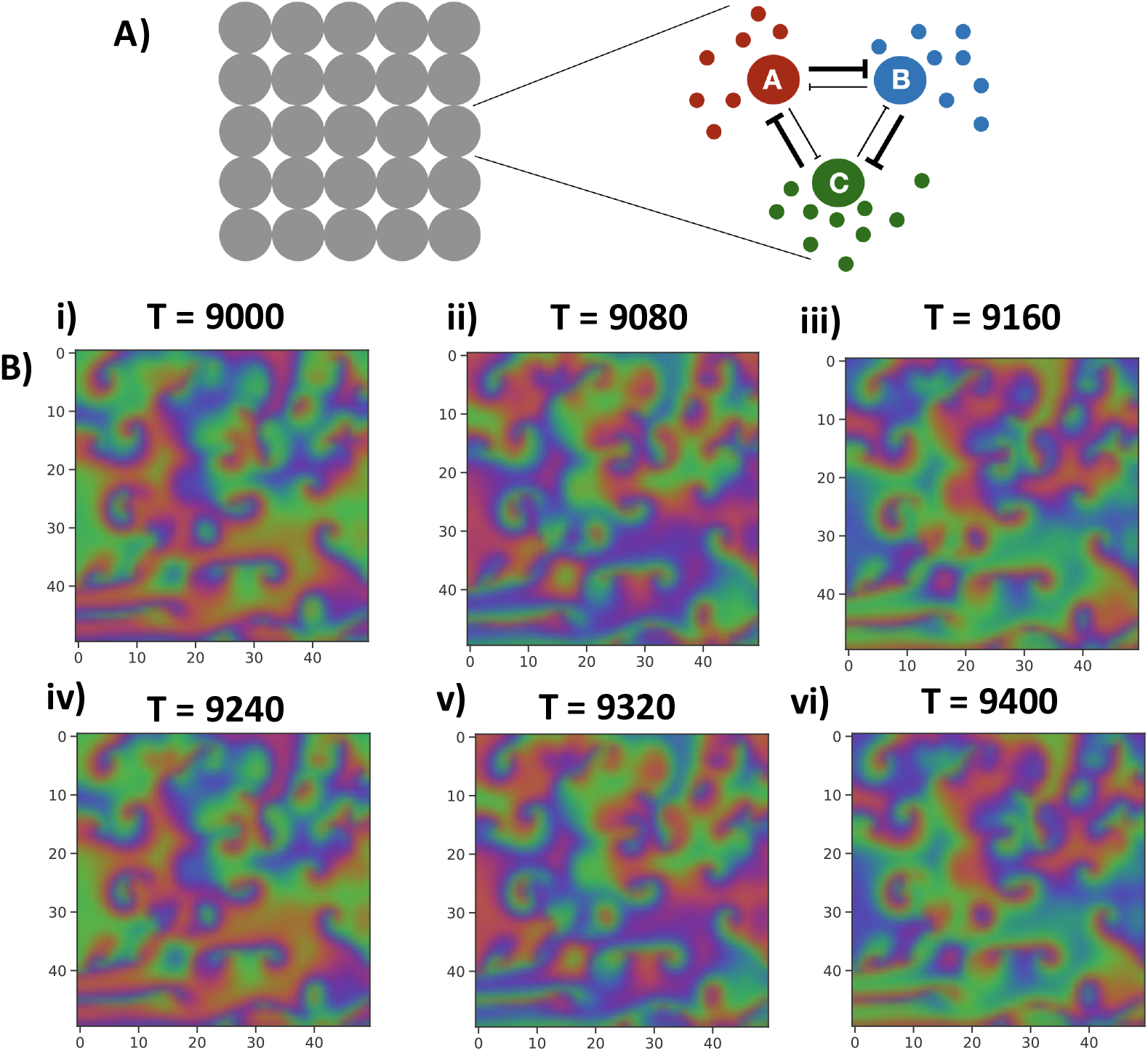
Repressilator as a special case of toggle triad: A) Schematic of Repressilator as a special case of toggle triad of three species in a cell coupled with the diffusion system. One cycle of the repression is weak as compared to the other one. B) An RGB (Red Green Blue) plot of levels of A, B and C respectively as a function of space at six different time-points: i) T = 9000 ii) T = 9080 iii) T = 9160 iv) T = 9240 v) T = 9320 vi) T = 9400.

We observe interesting dynamics when we couple the repressilator motif with the diffusion dynamics. Initially the system develops dynamic patterns which don’t repeat themselves. But after a time point (around T = 8500 here), they start repeating themselves. Although the molecules diffuse throughout the space the pattern repeats itself every 240 time steps. We also observe the local formation of spirals around which the pattern repeats itself (Figure 6 B i)-vi)). A video of the oscillatory patterns is presented in the supplementary information S3.4. The overlay plot is shown in the main while the separate expression levels for the molecules of gene A, gene B and C for the same parameter set are shown in Fig. S11. The oscillatory behavior is shown from the analytical calculation and its graphical representation of the three eigenvalues (one positive, and two complex eigenvalues with negative real part) for with diffusion in two-dimensional space as a function of the wavenumber *k* in Fig. S12.

## 3 Discussion

Many biological phenomena result from a constant tug-of-war between the contrasting aspects of stochastic and deterministic behavior, i.e. random fluctuations in gene expression, concentration of numerous intra-cellular biomolecules and their interactions via different signaling pathways, cell-to-cell variability, inter-cellular communication etc. ultimately result in some coordinated emergent phenomena of cellular division, differentiation and development [49]. Similarly, the biological systems which fall into the category of reaction-diffusion systems, biochemical “reactions” is a stabilizing/homogenizing/equilibrating process while molecular “diffusion” is trying to destabilize it. These two contradictory processes results in the spontaneous emergence of spatiotemporal patterns. “Reaction” and “Diffusion” might refer to intra-cellular biochemical reactions and inter-cellular signaling/communication (at the tissue-level) respectively. Unraveling the underlying mechanisms of these processes is the key challenging problem in synthetic [28] and developmental biology. An asymptotically stable homogeneous steady state of an inherently nonlinear system becoming unstable by the effect of self- and/or cross-diffusion is known as diffusion-driven instability or Turing instability [50].

Spatiotemporal patterns have been studied in multifarious contexts with different associated length-scales and time-scales. In population ecology - the systems of interacting population of prey-dependent predator–prey models with the influence of self- and cross-diffusion, gestation delays, weak Allele effects, analyzing formation of different patterns is an interesting domain of study [51–54]. Bacterial pattern formation [55] has been studied to understand the effects of the biofilms and quorum-sensing in the wild-type and mutant colonies of the pathogenic bacteria *Staphylococcus aureus* [56], and *Escherichia coli* [57, 58] by simulating the gene regulatory systems.

Many theoretical [59–63] and some experimental [64–66] studies suggest that Reaction-Diffusion mechanism plays the key role in bringing about various patterns (stripes, spots, patches, color variations) on animals, birds, insects or fish. Using microscopy and fluorescence in-situ hybridization experimental techniques to know more about gene locus and nuclear architecture, Fritsch *et al*. [67] observed beautiful expression patterns of gap and pair-rule genes such as hunchback, bicoid in Drosophila embryo. Reaction-Diffusion model of hunchback transcription, and bicoid self-regulation in bicoid gradient during embryogenesis of Drosophila exhibits spatial bistability [2].

Barbier *et al*. constructed synthetic toggle switches in *E. coli* and kept them under the influence of diffusive gradients of an external inducer and a regulator and observed, characterized, and tuned parametrically the robust spatiotemporal patterning exhibiting bistability and hysteresis [32]. Besides naturally existing patterns, synthetic patterns have been engineered in microbial communities [55, 68]. Theoretical explorations of spatial patterns (transient but persistent) formed from quorum-sensing modules of bistable toggle switch and a four-state GRN with diffusible molecules in one- and two-dimensional space, having leaky gene expression and cross-talk [57] is also worth mentioning. Recently Tompkins *et al*. performed quantitative analysis of Turing’s hypothesis and experimental testing of it on an emulsion of aqueous droplets (mimicking circular array of identical biological cells) [69]. This also opens up applications of Turing model in material science. Though there have been a plethora of theoretical explorations on biological pattern formation over decades, but studies of these in the context of gene regulatory networks is a very recent paradigm of interest.

We have presented a detailed analysis of spatiotemporal patterns emerging from the spatially cued molecules involved in two- and three-node regulatory networks showing multistable or oscillatory behavior. The analysis begins with the simple two-node network of toggle switch, having both monostable and bistable behavior. Toggle switch coupled with self loops adds an additional stable state. Toggle Triad is another example which shows all the mono-, bi- and tristable behavior in the spatiotemporal dynamics. Repressilator exhibits an oscillatory behavior. Full nonlinear model should be solved to gain more precise and detailed insights instead of Linear Stability Analysis. The latter captures majority of the dynamical features of the system qualitatively, leaving out the intricate details. We have developed a minimal, generic and coarse-grained but strong theory which has the potential to capture various spatiotemporal patterns taking into account the structure and different multistable parameter regimes of the underlying gene regulatory networks.

In simulations, we assumed free diffusion across cell membranes. In real biological systems, the positional and structural information about the cell-membranes and other membrane-bound organelles is required to set up the diffusion gradient of the molecules. Theoretically, diverse patterns can be generated from partial differential equations which falls into the broad category of the Reaction-Diffusion systems. Visually similar *in-silico* patterns can be generated from equations having different reaction and diffusion terms. Thus, an interplay of more than one factor can occur to bring about a biological stationary pattern.

Animals can have different body structures, sizes, shapes and curvatures which demands tackling of the nonlinear multivariate system of partial differential equations to be solved for different geometries/topologies - spherical, cylindrical etc. For simplicity, we have taken diffusion coefficient for different genes to be the same. Realistically, these should differ inherently and along different directions and might be a function of space due to anisotropy in diffusion of molecules. Thus, next steps include incorporating these realistic features and different initial and boundary conditions imitating more realistic biological scenarios. The systems can be studied for more specific biological cases where we can tune parameters using experimental results as inputs. In our analysis, we have diagonal diffusivity tensors, as we have only self-diffusion of the biomolecules. This assumption is reasonable because molecular diffusion of one molecule does not depend directly necessarily on others. But we can extend these scenarios and tackle more realistic cases which will have non-zero off-diagonal elements in the diffusivity tensor to capture the cross-diffusion dependent on the concentration gradient of some external component (inducer and/or regulator). Here, we have analyzed bivariate (toggle switch) and trivariate systems (Toggle Triad and Repressilator). In principle, systems can be in general be multivariate (toggle polygons and other networks) described by multiple nonlinear coupled partial differential equations. The essence of the problem remains the same while the complexity of solving it increases manifold due to a larger number of parameters.

Despite providing valuable insights into the emergent dynamics of pattern formation, our analysis here has multiple limitations. First, we have modeled Reaction-Diffusion systems, i.e. added a diffusive term (diffusion coefficient multiplied by the net flux (second-order partial differentiation with respect to space) of the (concentration of) particular gene, provided there are no other source or sink within the space) in the PDEs describing the evolutionary dynamics of the motifs. We do not consider the length and time scales explicitly in our problem, rather these scales are implicitly present in the diffusion coefficient term. Contrastingly, considering diffusion-limited biomolecular reaction kinetics (rate of reactions being functions of the diffusive transport of molecules) is important in some scenarios [70]. Second, we have considered a static lattice with fixed domain in our simulations. Cell division and death (apoptosis/necrosis) is not relevant in the timescales we are considering. Third, transcription factors are not known to diffuse across cell membranes. Thus, the molecules considered here are not necessarily transcription factors, but can represent at a coarse-grained level downstream signaling/regulatory molecule that can diffuse at a larger length-scale.

## 4 Methods and Materials

### 4.1 Analytical formalism

We have studied few two- and three-gene regulatory network motifs which exhibit inherent multistable and oscillatory behaviors, viz., genetic toggle switch, genetic toggle switch coupled with double self-activation, toggle triad, repressilator. We have written down the sets of partial differential equations representing the Reaction-Diffusion systems for each of these motifs. Numerically solving the Reaction systems (without including the term for the diffusing molecules), we first find the sets of real solutions (coordinates) for the stable fixed points/saddle nodes of the multistable/oscillatory states of the motifs. Linearizing around the stable fixed points/saddle modes, we obtain the Linear Stability Matrix 𝒮. Solving the matrix 𝒮, we find the traces, determinants and eigenvalues for the systems, the signs and types of which represents the stability of the systems (refer to text of SI for the detailed analytical treatment for each motif). We follow the same analytical treatment with molecular diffusion in one- and two-dimensional space. The resultant trace, determinant and eigenvalues of the reaction-diffusion systems (diffusive GRNs) are functions of the wavenumber *k* (see text in SI for details). To make the distinctions clear and comprehensible, the detailed analytical treatment for the Reaction systems and Reaction-Diffusion systems have been done (see SI text), separately for all the GRNs.

### 4.2 Simulation details

#### 4.2.1 Temporal dynamics of the GRNs

To solve the coupled ordinary differential equations (ODEs) (equations written in SI) of the network motif interactions, we integrate the ODESs using the MATLAB ode45 solver. We solve the ODEs for 100 initial conditions chosen from the range [0, 1.5 * *g*_*A*_*/γ*_*A*_] for total time of 200 units.

#### 4.2.2 Bifurcation Diagram

To calculate and plot the bifurcation diagrams of the coupled ODEs, we used MATCONT (a bifurcation analysis tool in MATLAB).

#### 4.2.3 Phase Diagram

We first divided the phase space in 50 × 50 cells. Then, for each cell, we calculated 50 trajectories, each starting with different initial conditions. We then calculate the number of states from these trajectories and plot the heatmap of the phase space.

#### 4.2.4 Spatiotemporal dynamics in one dimension

Consider a one dimensional chain of cells, each having a toggle switch with diffusing molecules. To solve the reaction diffusion system in 1 dimension we used the pdepe solver in MATLAB. We solve the system for 1D array of 400 cells with Neumann boundary condition.

#### 4.2.5 Spatiotemporal dynamics in two dimension

Now, we consider a two dimensional square lattice of population of cells, each having a toggle switch with diffusing molecules. To solve this system, we wrote a solver using central difference in space and forward difference in time (in python). We divided the space into lattice with 100×100 points and used Neumann boundary conditions. The initial levels of the proteins is chosen from the range [0, 1.5 * *g*_*A*_*/γ*_*A*_].

#### 4.2.6 Data Availability Statement

All codes are publicly available on the GitHub page of D.S. (https://github.com/Divyoj-Singh/Spatial_patterning_GRNs).

## 5 Author Information

### Corresponding Authors

Ushasi Roy

Mohit K. Jolly

### Author Contributions

UR and MKJ planned and led the work. UR, DS, NV, CH carried out the research. UR and DS prepared the initial manuscript. UR, DS, and MKJ finalized the draft and made intellectual contributions. UR and DS contributed equally to the work, are co-first authors, and they can exchange the author-order in their CV. NV and CH contributed equally to the work, are co-second authors, and they can exchange the author-order in their CV.

### Notes

The authors declare no competing financial interest.

## Acknowledgement

UR acknowledges C. V. Raman Postdoctoral Fellowship from Indian Institute of Science, Bangalore, India. DS, NV and CH are supported by KVPY fellowship awarded by Department of Science and Technology (DST), Government of India. MKJ is supported by Ramanujan Fellowship (SB/S2/RJN 049/2018) awarded by the Science and Engineering Research Board (SERB), DST, Government of India.

## References

[1] Stephen DiNardo, Jill Heemskerk, Scott Dougan, and Patrick H O’Farrell. The making of a maggot: patterning the drosophila embryonic epidermis. Current opinion in genetics & development, 4(4):529–534, 1994.

[2] Francisco JP Lopes, Fernando MC Vieira, David M Holloway, Paulo M Bisch, and Alexander V Spirov. Spatial bistability generates hunchback expression sharpness in the drosophila embryo. PLoS computational biology, 4(9):e1000184, 2008.

[3] Edgar Herrera-Delgado, Ruben Perez-Carrasco, James Briscoe, and Peter Sollich. Memory functions reveal structural properties of gene regulatory networks. PLoS computational biology, 14(2):e1006003, 2018.

[4] Bard Ermentrout. Neural networks as spatio-temporal pattern-forming systems. Reports on progress in physics, 61(4):353, 1998.

[5] Ryo Futahashi, Hiroko Shirataki, Takanori Narita, Kazuei Mita, and Haruhiko Fujiwara. Comprehensive microarray-based analysis for stage-specific larval camouflage pattern-associated genes in the swallowtail butterfly, papilio xuthus. BMC biology, 10(1):1–22, 2012.

[6] Hans Meinhardt. A model for pattern formation of hypostome, tentacles, and foot in hydra: how to form structures close to each other, how to form them at a distance. Developmental biology, 157(2):321–333, 1993.

[7] James Briscoe and Stephen Small. Morphogen rules: design principles of gradient-mediated embryo patterning. Development, 142:3996–4009, 12 2015.

[8] Peter A. Lawrence and Gary Struhl. Morphogens, compartments, and pattern: Lessons from drosophila? Cell, 85:951–961, 6 1996.

[9] Melinda Liu and Perkins Id. Implications of diffusion and time-varying morphogen gradients for the dynamic positioning and precision of bistable gene expression boundaries. PLOS Computational Biology, 17:e1008589, 6 2021.

[10] Alan Mathison Turing. The chemical basis of morphogenesis. Bulletin of mathematical biology, 52(1):153–197, 1990.

[11] AN Zaikin and AM Zhabotinsky. Concentration wave propagation in two-dimensional liquid-phase self-oscillating system. Nature, 225(5232):535–537, 1970.

[12] Arthur T Winfree. Spiral waves of chemical activity. Science, 175(4022):634–636, 1972.

[13] Alfred Gierer and Hans Meinhardt. A theory of biological pattern formation. Kybernetik, 12(1):30–39, 1972.

[14] Vincent Castets, Etiennette Dulos, Jacques Boissonade, and Patrick De Kepper. Experimental evidence of a sustained standing turing-type nonequilibrium chemical pattern. Physical review letters, 64(24):2953, 1990.

[15] P De Kepper, V Castets, E Dulos, and J Boissonade. Turing-type chemical patterns in the chlorite-iodide-malonic acid reaction. Physica D: Nonlinear Phenomena, 49(1-2):161–169, 1991.

[16] Adolfo Sanz-Anchelergues, Anatol M Zhabotinsky, Irving R Epstein, and Alberto P Munuzuri. Turing pattern formation induced by spatially correlated noise. Physical Review E, 63(5):056124, 2001.

[17] Hans G Othmer and LE Scriven. Instability and dynamic pattern in cellular networks. Journal of theoretical biology, 32(3):507–537, 1971.

[18] Istvan Lengyel and Irving R Epstein. A chemical approach to designing turing patterns in reaction-diffusion systems. Proceedings of the National Academy of Sciences, 89(9):3977–3979, 1992.

[19] Mark C Cross and Pierre C Hohenberg. Pattern formation outside of equilibrium. Reviews of modern physics, 65(3):851, 1993.

[20] Justin Hsia, William J Holtz, Daniel C Huang, Murat Arcak, and Michel M Maharbiz. A feedback quenched oscillator produces turing patterning with one diffuser. PLoS computational biology, 8(1):e1002331, 2012.

[21] Hiroki Miyazako, Yutaka Hori, and Shinji Hara. Turing instability in reaction-diffusion systems with a single diffuser: characterization based on root locus. In 52nd IEEE Conference on Decision and Control, pages 2671–2676. IEEE, 2013.

[22] Stephen Payne, Bochong Li, Yangxiaolu Cao, David Schaeffer, Marc D Ryser, and Lingchong You. Temporal control of self-organized pattern formation without morphogen gradients in bacteria. Molecular systems biology, 9(1):697, 2013.

[23] Yangxiaolu Cao, Marc D Ryser, Stephen Payne, Bochong Li, Christopher V Rao, and Lingchong You. Collective space-sensing coordinates pattern scaling in engineered bacteria. Cell, 165(3):620–630, 2016.

[24] Daniel J Cohen, Roberto C Morfino, and Michel M Maharbiz. A modified consumer inkjet for spatiotemporal control of gene expression. PloS one, 4(9):e7086, 2009.

[25] Takayuki Sohka, Richard A Heins, and Marc Ostermeier. Morphogen-defined patterning of escherichia coli enabled by an externally tunable band-pass filter. Journal of biological engineering, 3(1):1–9, 2009.

[26] Subhayu Basu, Rishabh Mehreja, Stephan Thiberge, Ming-Tang Chen, and Ron Weiss. Spatiotemporal control of gene expression with pulse-generating networks. Proceedings of the National Academy of Sciences, 101(17):6355–6360, 2004.

[27] Subhayu Basu, Yoram Gerchman, Cynthia H Collins, Frances H Arnold, and Ron Weiss. A synthetic multicellular system for programmed pattern formation. Nature, 434(7037):1130–1134, 2005.

[28] Nan Luo, Shangying Wang, and Lingchong You. Synthetic pattern formation. Biochemistry, 58(11):1478–1483, 2019.

[29] Jeroen S van Zon, Marco J Morelli, Sorin Tânase-Nicola, and Pieter Rein ten Wolde. Diffusion of transcription factors can drastically enhance the noise in gene expression. Biophysical journal, 91(12):4350–4367, 2006.

[30] Gašper Tkačik and William Bialek. Diffusion, dimensionality, and noise in transcriptional regulation. Physical Review E, 79(5):051901, 2009.

[31] David Cottrell, Peter S Swain, and Paul F Tupper. Stochastic branching-diffusion models for gene expression. Proceedings of the National Academy of Sciences, 109(25):9699–9704, 2012.

[32] Içvara Barbier, Rubén Perez-Carrasco, and Yolanda Schaerli. Controlling spatiotemporal pattern formation in a concentration gradient with a synthetic toggle switch. Molecular systems biology, 16(6):e9361, 2020.

[33] Ruben Perez-Carrasco, Chris P Barnes, Yolanda Schaerli, Mark Isalan, James Briscoe, and Karen M Page. Combining a toggle switch and a repressilator within the ac-dc circuit generates distinct dynamical behaviors. Cell systems, 6(4):521–530, 2018.

[34] Luis Diambra, Vivek Raj Senthivel, Diego Barcena Menendez, and Mark Isalan. Cooperativity to increase turing pattern space for synthetic biology. ACS synthetic biology, 4(2):177–186, 2015.

[35] Timothy S Gardner, Charles R Cantor, and James J Collins. Construction of a genetic toggle switch in escherichia coli. Nature, 403(6767):339–342, 2000.

[36] Dongya Jia, Mohit Kumar Jolly, William Harrison, Marcelo Boareto, Eshel Ben-Jacob, and Herbert Levine. Operating principles of tristable circuits regulating cellular differentiation. Physical biology, 14(3):035007, 2017.

[37] Atchuta Srinivas Duddu, Sarthak Sahoo, Souvadra Hati, Siddharth Jhunjhunwala, and Mohit Kumar Jolly. Multi-stability in cellular differentiation enabled by a network of three mutually repressing master regulators. Journal of the Royal Society Interface, 17(170):20200631, 2020.

[38] Michael B Elowitz and Stanislas Leibler. A synthetic oscillatory network of transcriptional regulators. Nature, 403(6767):335–338, 2000.

[39] Joseph X Zhou and Sui Huang. Understanding gene circuits at cell-fate branch points for rational cell reprogramming. Trends in genetics, 27(2):55–62, 2011.

[40] Francisco JP Lopes, Alexander V Spirov, and Paulo M Bisch. The role of bicoid cooperative binding in the patterning of sharp borders in drosophila melanogaster. Developmental biology, 370(2):165–172, 2012.

[41] Oleg A Igoshin, Rui Alves, and Michael A Savageau. Hysteretic and graded responses in bacterial two-component signal transduction. Molecular microbiology, 68(5):1196–1215, 2008.

[42] Ophelia S Venturelli, Hana El-Samad, and Richard M Murray. Synergistic dual positive feedback loops established by molecular sequestration generate robust bimodal response. Proceedings of the National Academy of Sciences, 109(48):E3324–E3333, 2012.

[43] Ophelia S Venturelli, Ignacio Zuleta, Richard M Murray, and Hana El-Samad. Population diversification in a yeast metabolic program promotes anticipation of environmental shifts. PLoS biology, 13(1):e1002042, 2015.

[44] Mingyang Lu, Mohit Kumar Jolly, Herbert Levine, José N. Onuchic, and Eshel Ben-Jacob. MicroRNA-based regulation of epithelial-hybrid-mesenchymal fate determination. Proceedings of the National Academy of Sciences of the United States of America, 110(45):18144–18149, nov 2013.

[45] Kishore Hari, Burhanuddin Sabuwala, Balaram Vishnu Subramani, Caterina AM La Porta, Stefano Zapperi, Francesc Font-Clos, and Mohit Kumar Jolly. Identifying inhibitors of epithelial–mesenchymal plasticity using a network topology-based approach. NPJ systems biology and applications, 6(1):1–12, 2020.

[46] Mingyang Lu, Mohit Kumar Jolly, Ryan Gomoto, Bin Huang, Jose Onuchic, and Eshel Ben-Jacob. Tristability in cancer-associated microrna-tf chimera toggle switch. The journal of physical chemistry B, 117(42):13164–13174, 2013.

[47] Shubham Tripathi, David A. Kessler, and Herbert Levine. Biological networks regulating cell fate choice are minimally frustrated. Phys. Rev. Lett., 125:088101, Aug 2020.

[48] Béla Novák and John J Tyson. Design principles of biochemical oscillators. Nature reviews Molecular cell biology, 9(12):981–991, 2008.

[49] Arjun Raj and Alexander Van Oudenaarden. Nature, nurture, or chance: stochastic gene expression and its consequences. Cell, 135(2):216–226, 2008.

[50] Syed Shahed Riaz and Deb Shankar Ray. Diffusion and mobility driven instabilities in a reaction-diffusion system: a review. In INDIAN JOURNAL OF PHYSICS AND PROCEEDINGS OF THE INDIAN ASSOCIATION FOR THE CULTIVATION OF SCIENCE-NEW SERIES-, volume 81, page 1177. NOT KNOWN, 2007.

[51] Lakshmi Narayan Guin, Mainul Haque, and Prashanta Kumar Mandal. The spatial patterns through diffusion-driven instability in a predator–prey model. Applied Mathematical Modelling, 36(5):1825–1841, 2012.

[52] Yexuan Li, Hua Liu, Yumei Wei, and Ming Ma. Turing pattern of a reaction-diffusion predator-prey model with weak allee effect and delay. In Journal of Physics: Conference Series, volume 1707, page 012025. IOP Publishing, 2020.

[53] Ye Xuan Li, Hua Liu, Yu Mei Wei, Ming Ma, Gang Ma, and Jing Yan Ma. Population dynamic study of prey-predator interactions with weak allee effect, fear effect, and delay. Journal of Mathematics, 2022, 2022.

[54] Jordi van Gestel, Tasneem Bareia, Bar Tenennbaum, Alma Dal Co, Polina Guler, Nitzan Aframian, Shani Puyesky, Ilana Grinberg, Glen G. D’Souza, Zohar Erez, Martin Ackermann, and Avigdor Eldar. Short-range quorum sensing controls horizontal gene transfer at micron scale in bacterial communities. Nature Communications 2021 12:1, 12:1–11, 4 2021.

[55] Salva Duran-Nebreda, Jordi Pla, Blai Vidiella, Jordi Piñero, Nuria Conde-Pueyo, and Ricard Solé. Synthetic lateral inhibition in periodic pattern forming microbial colonies. ACS Synthetic Biology, 10:277–285, 2 2021.

[56] A Oelker, T Horger, and C Kuttler. From staphylococcus aureus gene regulation to its pattern formation. Journal of Mathematical Biology, 78(7):2207–2234, 2019.

[57] Marcella M Gomez and Murat Arcak. A tug-of-war mechanism for pattern formation in a genetic network. ACS synthetic biology, 6(11):2056–2066, 2017.

[58] Alma Dal Co, Simon van Vliet, Daniel Johannes Kiviet, Susan Schlegel, and Martin Ackermann. Short-range interactions govern the dynamics and functions of microbial communities. Nature Ecology & Evolution 2020 4:3, 4:366–375, 2 2020.

[59] Shigeru Kondo. The reaction-diffusion system: a mechanism for autonomous pattern formation in the animal skin. Genes to Cells, 7(6):535–541, 2002.

[60] J D Murray. A pre-pattern formation mechanism for animal coat markings. Journal of Theoretical Biology, 88(1):161–199, 1981.

[61] Cheng-Ming Chuong and Michael K Richardson. Pattern formation today. The International journal of developmental biology, 53(5-6):653, 2009.

[62] Hans Meinhardt. Models of biological pattern formation. New York, 118, 1982.

[63] Hans Meinhardt. The algorithmic beauty of sea shells. Springer Science & Business Media, 2009.

[64] Patricia Macauley Bode and Hans R Bode. Formation of pattern in regenerating tissue pieces of hydra attenuata: Ii. degree of proportion regulation is less in the hypostome and tentacle zone than in the tentacles and basal disc. Developmental biology, 103(2):304–312, 1984.

[65] Kemble Yates and Edward Pate. A cascading development model for amphibian embryos. Bulletin of mathematical biology, 51(5):549–578, 1989.

[66] Yoram Schiffmann. An hypothesis: phosphorylation fields as the source of positional information and cell differentiation—(camp, atp) as the universal morphogenetic turing couple. Progress in biophysics and molecular biology, 56(2):79–105, 1991.

[67] Cornelia Fritsch, Ginette Ploeger, and Donna J Arndt-Jovin. Drosophila under the lens: imaging from chromosomes to whole embryos. Chromosome Research, 14(4):451–464, 2006.

[68] Içvara Barbier, Hadiastri Kusumawardhani, and Yolanda Schaerli. Engineering synthetic spatial patterns in microbial populations and communities. 2022.

[69] Nathan Tompkins, Ning Li, Camille Girabawe, Michael Heymann, G Bard Ermentrout, Irving R Epstein, and Seth Fraden. Testing turing’s theory of morphogenesis in chemical cells. Proceedings of the National Academy of Sciences, 111(12):4397–4402, 2014.

[70] Sumantra Sarkar. Concentration dependence of diffusion-limited reaction rates and its consequences. Physical Review X, 10(4):041032, 2020.

