## Supplementary Figures/Information for "Spatiotemporal Patterning enabled by Gene Regulatory Networks"

### Spatial Patterns enabled by Gene Regulatory Networks Supplementary Information

#### 1 Analytical formulation

Here, we present a detailed description of how we might analyze a Reaction-Diffusion system analytically, in general. We begin our discussion with two-component Reaction systems. We illustrate how to formulate specific mathematical quantities like Trace, Determinant and Eigenvalues fom the Linear Stability Matrix to analyze the stabilities of the systems for four specific examples of inherently multistable and oscilatory gene regulatory motifs - Toggle Switch (TS), Toggle Switch with self-activation (TSSA), Toggle Triad (TT) and Reressilator (same as in main text). Then we discuss about Reaction-Diffusion systems. Lastly we illustrate how the incorporation of one-dimensional and two-dimensional diffusion changes the mathematical expressions of the Reaction Systems for these specific cases. The systems without diffusion, and with diffusion in one- and two- spatial dimensions are depicted graphically in the main and supplementary, wherever relevant.

##### 1.1 Reaction Systems

These are often called well-mixed systems (no diffusion). There exists spatially uniform steady state(s), stable to perturbations.

$$\begin{aligned} \frac{dA}{dt} &= f(A, B) \\ \frac{dB}{dt} &= g(A, B) \end{aligned} \tag{1}$$

Let $(A_0, B_0)$ be solution (coordinates of the equilibrium point) at the steady-state. This is a multi-dimensional and an inherently nonlinear system, hard to analyze analytically. So, for analytical tractability, let's linearize $f(A, B)$ and $g(A, B)$ around $(A_0, B_0)$. Linearization (taking Taylor expansion and retaining only first order terms) gives,

$$\begin{aligned} f(A, B) &\approx f(A_0, B_0) + f_A\delta_A + f_B\delta_B \\ g(A, B) &\approx g(A_0, B_0) + g_A\delta_A + g_B\delta_B \end{aligned}$$

where $f_A$, $f_B$, $g_A$ and $g_B$ are the first derivatives evaluated at $(A_0, B_0)$

$$
\begin{aligned}
f_A = \left.\frac{\partial f}{\partial A}\right|_{(A_0,B_0)}, \quad f_B = \left.\frac{\partial f}{\partial B}\right|_{(A_0,B_0)} \\
g_A = \left.\frac{\partial g}{\partial A}\right|_{(A_0,B_0)}, \quad g_B = \left.\frac{\partial g}{\partial B}\right|_{(A_0,B_0)}
\end{aligned} \tag{2}
$$

and $\delta A(t)$ and $\delta B(t)$ are the perturbations from the steady state $(A_0, B_0)$,

$$
\begin{aligned}
\delta A(t) &= A(t) - A_0 \\
\delta B(t) &= B(t) - B_0.
\end{aligned}
$$

Thus the linearized system is expressed as

$$
\begin{aligned}
\frac{d}{dt}\delta A &= f_A \delta A + f_B \delta B \\
\frac{d}{dt}\delta B &= g_A \delta A + g_B \delta B
\end{aligned}
$$

or in a compact form

$$
\frac{d}{dt}\begin{bmatrix}\delta A \\ \delta B\end{bmatrix} = \begin{bmatrix} f_A & f_B \\ g_A & g_B \end{bmatrix}\begin{bmatrix}\delta A \\ \delta B\end{bmatrix} = \mathcal{S}\begin{bmatrix}\delta A \\ \delta B\end{bmatrix} \tag{3}
$$

where $\mathcal{S}$ is the linear stability matrix. Let us assume the solution of this linear system to be of the form

$$
\begin{bmatrix}\delta A \\ \delta B\end{bmatrix} = e^{\lambda t}\begin{bmatrix}\delta A_0 \\ \delta B_0\end{bmatrix}
$$

where $\delta A_0$ and $\delta B_0$ are constants. The trace $\tau$ and determinant $\Delta$ of the linear stability matrix $\mathcal{S}$ is given by

$$
\begin{aligned}
\tau &= Tr(\mathcal{S}) = f_A + g_B \\
\Delta &= Det(\mathcal{S}) = f_A g_B - f_B g_A
\end{aligned} \tag{4}
$$

The characteristic equation, expressed in terms of the trace and determinant, is given by

$$
\lambda^2 - \tau\lambda + \Delta = 0
$$

where $\lambda$s are the eigenvalues of the matrix $\mathcal{S}$. Explicitly,

$$
\lambda_{\pm} = \frac{1}{2}\left[\tau \pm \sqrt{\tau^2 - 4\Delta}\right] \tag{5}
$$

Stability of the system, i.e., $Re(\lambda) < 0$, is guaranteed when the following are satisfied

$$
\begin{aligned}
\tau &= Tr(\mathcal{S}) = f_A + g_B < 0 \\
\Delta &= Det(\mathcal{S}) = f_A g_B - f_B g_A > 0
\end{aligned} \tag{6}
$$

If we have multiple steady states for a system, we need to analyze and choose the stable one.

#### 1.2 Toggle Switch

The governing equation for genetic toggle switch (TS) - two genes mutually repressing each other - in terms of the shifted Hill function $\mathcal{H}$ is given by

$$\begin{aligned}\frac{dA}{dt} &= f(A,B) = g_A\mathcal{H}(B, B_0A, n_{BA}, l_{BA}) - \gamma_A A \\ \frac{dB}{dt} &= g(A,B) = g_B\mathcal{H}(A, A_0B, n_{AB}, l_{AB}) - \gamma_B B\end{aligned} \tag{7}$$

where $g_A$ and $g_B$ are the production rates, $\gamma_A$ and $\gamma_B$ are the degradation rates and $\mathcal{H}(B, B_0A, n_{BA}, l_{BA})$ and $\mathcal{H}(A, A_0B, n_{AB}, l_{AB})$ are the shifted Hill functions given by

$$\begin{aligned}\mathcal{H}(B, B_0A, n_{BA}, l_{BA}) &= H_-(B, B_0A, n_{BA}, l_{BA}) + l_{BA}H_+(B, B_0A, n_{BA}, l_{BA}) \\ &= \frac{B_0A^{n_{BA}}}{B_0A^{n_{BA}} + B^{n_{BA}}} + l_{BA}\frac{B^{n_{BA}}}{B_0A^{n_{BA}} + B^{n_{BA}}} \\ \mathcal{H}(A, A_0B, n_{AB}, l_{AB}) &= H_-(A, A_0B, n_{AB}, l_{AB}) + l_{AB}H_+(A, A_0B, n_{AB}, l_{AB}) \\ &= \frac{A_0B^{n_{AB}}}{A_0B^{n_{AB}} + A^{n_{AB}}} + l_{AB}\frac{A^{n_{AB}}}{A_0B^{n_{AB}} + A^{n_{AB}}}\end{aligned} \tag{8}$$

where $B_0A$ and $A_0B$ are the thresholds, $n_{BA}$ and $n_{AB}$ are the Hill coefficients, $l_{BA}$ and $l_{AB}$ are the fold changes. $H_-$ and $H_+$ denotes the repression and activation from the basal expression level. Now the first derivatives of the reaction terms are as follows

$$\begin{aligned}f_A &= \left.\frac{\partial f(A,B)}{\partial A}\right|_{(A_0,B_0)} = -\gamma_A \\ f_B &= \left.\frac{\partial f(A,B)}{\partial v}\right|_{(A_0,B_0)} = \frac{g_A n_{BA}(l_{BA}-1)B_0A^{n_{BA}}v^{(n_{BA}-1)}}{(B_0A^{n_{BA}} + v^{n_{BA}})^2} \\ g_A &= \left.\frac{\partial g(A,B)}{\partial A}\right|_{(A_0,B_0)} = \frac{g_B n_{AB}(l_{AB}-1)A_0B^{n_{AB}}u^{(n_{AB}-1)}}{(A_0B^{n_{AB}} + A^{n_{AB}})^2} \\ g_B &= \left.\frac{\partial g(A,B)}{\partial v}\right|_{(A_0,B_0)} = -\gamma_B\end{aligned} \tag{9}$$

Linear Stability Matrix $\mathcal{S}$, Trace $\tau$, Determinant $\Delta$, and the Eigenvalues $\lambda_\pm$ are given by Equations (3, 4, 5) respectively, all of which are combinations of $f_A$, $f_B$, $g_A$ and $g_B$ as expressed above in Eq.(9). This motif results in both monostable as well as bistable behavior for different values of the parameters (refer to Fig. 1 in main text).

#### 1.3 Toggle Switch with double self-activation

The governing equations for genetic toggle switch with double self-activation (TSSA) (discussed in detail in Sec. 3 and the behavior shown in Fig. 3 of main text) are

$$
\begin{aligned}
\frac{dA}{dt} &= g_A \mathcal{H}(B, B_0 A, n_{BA}, l_{BA}) \mathcal{H}(B, A_0 A, n_{AA}, l_{AA}) - \gamma_A A \\
\frac{dB}{dt} &= g_B \mathcal{H}(A, A_0 B, n_{AB}, l_{AB}) \mathcal{H}(B, B_0 B, n_{BB}, l_{BB}) - \gamma_B B
\end{aligned} \qquad (10)
$$

where $\mathcal{H}(B, B_0 A, n_{BA}, l_{BA})$ and $\mathcal{H}(A, A_0 B, n_{AB}, l_{AB})$ are the shifted Hill functions as given in Eq.(8) in the previous section, while the other two shifted Hill functions are for the self-activation loops, given by

$$
\begin{aligned}
\mathcal{H}(B, A_0 A, n_{AA}, l_{AA}) &= H_{-}(B, A_0 A, n_{AA}, l_{AA}) + l_{AA} H_{+}(B, A_0 A, n_{AA}, l_{AA}) \\
&= \frac{A_0 A^{n_{AA}}}{A_0 A^{n_{AA}} + B^{n_{AA}}} + l_{AA} \frac{B^{n_{AA}}}{A_0 A^{n_{AA}} + B^{n_{AA}}} \\
\mathcal{H}(A, B_0 B, n_{BB}, l_{BB}) &= H_{-}(A, B_0 B, n_{BB}, l_{BB}) + l_{BB} H_{+}(A, B_0 B, n_{BB}, l_{BB}) \\
&= \frac{B_0 B^{n_{BB}}}{B_0 B^{n_{BB}} + A^{n_{BB}}} + l_{BB} \frac{A^{n_{BB}}}{B_0 B^{n_{BB}} + A^{n_{BB}}}
\end{aligned} \qquad (11)
$$

The first derivatives of the reaction terms are given by

$$
\begin{aligned}
f_A &= \frac{-\gamma_A (A_0 A^{n_{AA}} + A^{n_{AA}})^2 (B_0 A^{n_{BA}} + B^{n_{BA}}) + g_A n_{AA} A_0 A^{n_{AA}} (l_{AA} - 1) A^{n_{AA}-1} (B_0 A^{n_{BA}} + l_{BA} B^{n_{BA}})}{(A_0 A^{n_{AA}} + A^{n_{AA}})^2 (B_0 A^{n_{BA}} + B^{n_{BA}})} \\
f_B &= \frac{g_A n_{BA} B_0 A^{n_{BA}} (l_{BA} - 1) v^{(n_{BA}-1)} (A_0 A^{n_{AA}} + l_{AA} u^{n_{AA}})}{(A_0 A^{n_{AA}} + A^{n_{AA}})(B_0 A^{n_{BA}} + B^{n_{BA}})^2} \\
g_A &= \frac{g_B n_{AB} A_0 B^{n_{AB}} (l_{AB} - 1) A^{(n_{AB}-1)} (B_0 B^{n_{BB}} + l_{BB} B^{n_{BB}})}{(B_0 B^{n_{BB}} + A^{n_{BB}})(A_0 B^{n_{AB}} + A^{n_{AB}})^2} \\
g_B &= \frac{-\gamma_B (B_0 B^{n_{BB}} + B^{n_{BB}})^2 (A_0 B^{n_{AB}} + A^{n_{AB}}) + g_B n_{BB} B_0 B^{n_{BB}} (l_{BB} - 1) B^{n_{BB}-1} (A_0 B^{n_{AB}} + l_{AB} A^{n_{AB}})}{(B_0 B^{n_{BB}} + B^{n_{BB}})^2 (A_0 B^{n_{AB}} + A^{n_{AB}})}
\end{aligned} \qquad (12)
$$

In addition to monostable and bistable behavior, similar to what we have observed in the the previous motif of simple Toggle Switch, TSSA shows tristable behavior also in some parameter regimes. Now, $f_A$, $f_B$, $g_A$ and $g_B$ (as in Eq.12) can be combined to get the expressions for the linear stability matrix $\mathcal{S}$, trace $\tau$, determinant $\Delta$ and eigenvalues $\lambda$, expressed in equations (**??**, **??**, **??**) can be determined and the stability of the system has been analyzed (see Fig. Fig. 3 in main text).

##### 1.4 Toggle Triad

The governing equations for Toggle Triad - three mutually coupled toggle switches (schematic shown in Fig. 4A of main text)

$$
\begin{aligned}
\frac{dA}{dt} &= f(A,B,C) = g_A \mathcal{H}(B, B_0 A, n_{BA}, l_{BA}) \mathcal{H}(C, C_0 A, n_{CA}, l_{CA}) - \gamma_A A \\
\frac{dB}{dt} &= g(A,B,C) = g_B \mathcal{H}(C, C_0 B, n_{CB}, l_{CB}) \mathcal{H}(A, A_0 B, n_{AB}, l_{AB}) - \gamma_B B \\
\frac{dC}{dt} &= h(A,B,C) = g_C \mathcal{H}(A, A_0 C, n_{AC}, l_{AC}) \mathcal{H}(B, B_0 C, n_{BC}, l_{BC}) - \gamma_C C \qquad (13)
\end{aligned}
$$

In the above equation, there are six different shifted Hill functions describing six different reactions, the exact functional form is easily deducible from Eq.(8) with appropriate substitutions. The first derivatives of the reaction terms are

$$
\begin{aligned}
f_A &= \frac{\partial f(A,B,C)}{\partial A} = -\gamma_A \\
f_B &= \frac{\partial f(A,B,C)}{\partial B} = \frac{g_A n_{BA} B_0 A^{n_{BA}} (l_{BA}-1) B^{(n_{BA}-1)} (C_0 A^{n_{CA}} + l_{CA} C^{n_{CA}})}{(B_0 A^{n_{BA}} + B^{n_{BA}})^2 (C_0 A^{n_{CA}} + C^{n_{CA}})} \\
f_C &= \frac{\partial f(A,B,C)}{\partial C} = \frac{g_A n_{CA} C_0 A^{n_{CA}} (l_{CA}-1) C^{(n_{CA}-1)} (B_0 A^{n_{BA}} + l_{BA} B^{n_{BA}})}{(B_0 A^{n_{BA}} + B^{n_{BA}}) (C_0 A^{n_{CA}} + C^{n_{CA}})^2} \qquad (14)
\end{aligned}
$$

$$
\begin{aligned}
g_A &= \frac{\partial g(A,B,C)}{\partial A} = \frac{g_B n_{AB} A_0 B^{n_{AB}} (l_{AB}-1) A^{(n_{AB}-1)} (C_0 B^{n_{CB}} + l_{CB} C^{n_{CB}})}{(A_0 B^{n_{AB}} + A^{n_{AB}})^2 (C_0 B^{n_{CB}} + C^{n_{CB}})} \\
g_B &= \frac{\partial g(A,B,C)}{\partial B} = -\gamma_B \\
g_C &= \frac{\partial g(A,B,C)}{\partial C} = \frac{g_B n_{CB} C_0 B^{n_{CB}} (l_{CB}-1) C^{(n_{CB}-1)} (A_0 B^{n_{AB}} + l_{AB} A^{n_{AB}})}{(A_0 B^{n_{AB}} + A^{n_{AB}}) (C_0 B^{n_{CB}} + C^{n_{CB}})^2} \qquad (15)
\end{aligned}
$$

$$
\begin{aligned}
h_A &= \frac{\partial h(A,B,C)}{\partial A} = \frac{g_C n_{AC} A_0 C^{n_{AC}} (l_{AC}-1) A^{(n_{AC}-1)} (B_0 C^{n_{BC}} + l_{BC} B^{n_{BC}})}{(A_0 C^{n_{AC}} + A^{n_{AC}})^2 (B_0 C^{n_{BC}} + B^{n_{BC}})} \\
h_B &= \frac{\partial h(A,B,C)}{\partial B} = \frac{g_C n_{BC} B_0 C^{n_{BC}} (l_{BC}-1) B^{(n_{BC}-1)} (A_0 C^{n_{AC}} + l_{AC} A^{n_{AC}})}{(A_0 C^{n_{AC}} + A^{n_{AC}}) (B_0 C^{n_{BC}} + B^{n_{BC}})^2} \\
h_C &= \frac{\partial h(A,B,C)}{\partial C} = -\gamma_C \qquad (16)
\end{aligned}
$$

This motif has three parameter regimes of stability, viz. monostability, bistability and tristability (refer to Fig. 4, 5 and 6 of main text). This is a three-component system. For these systems, the linear stability matrix $\mathcal{S}$, trace $\tau$, determinant $\Delta$ and eigenvalues $\lambda_{\pm}$ of $\mathcal{S}$ are given by the following

$$\begin{aligned}
\mathcal{S} &= \begin{bmatrix} f_A & f_B & f_c \\ g_A & g_B & g_c \\ h_A & h_B & h_c \end{bmatrix} \\
\tau &= f_A + f_B + f_C \\
\Delta &= f_A(g_B h_C - g_C h_B) + f_B(g_C h_A - g_A h_C) + f_C(g_A h_B - g_B h_A)
\end{aligned} \tag{17}$$

The eigenvalues $\lambda_{\pm}$ can be expressed as in Eq.5 where $\tau$ and $\Delta$ are substituted from the above equation (Eq.(17)).

#### 1.5 Repressilator - a limiting case of toggle triad

A simple repressilator is a motif formed from three genes, cyclically repressing one another. Hence there are three links (schematic shown in Fig. 7 in main text and Fig. S9 in SI). Now, we have six repressive connections in Toggle Triad (schematic in Fig. 4 in main text). In the equations for toggle triad 13, we consider the fold-changes $l_{BA}$, $l_{CB}$ and $l_{AC}$ to be large (close to 1) while the other three fold changes $l_{AB}$, $l_{BC}$ and $l_{CA}$, to be a very small (close to 0) positive quantities. Making three links weaker and three links stronger, we effectively achieve the configuration of the repressilator (schematic shown in Fig. S9). The stability of the system is explored in a similar way as Toggle Triad, just with a different set of parameters.

#### 1.6 Inclusion of DIFFUSION

Diffusion is basically a stabilizing process that makes the system homogeneous. But here we study diffusion of interacting chemicals/morphogens/genes/molecules and from the interaction of two stabilizing processes, instability could emerge, called diffusion-driven instability. Pattern is said to emerge or self-organize.

#### 1.7 Reaction-Diffusion Systems in 1D

In the non-mixed system (in presence of diffusion), the steady state becomes unstable because of diffusion. Such a system is often referred to as the Reaction-Diffusion (RD) systems. A RD system, for which we have assumed that there exists a temporally and spatially independent stationary solution, exhibits diffusion-driven instability, if the homogeneous steady state is stable to small perturbations in the absence of diffusion and the state eventually becomes unstable to small spatial perturbations when diffusion is present. So we assume that the main process driving the spatially inhomogeneous instability is diffusion and the underlying mechanism determines the spatial pattern that evolves. The governing equations for a 2-component (A,B) RD system in one-dimension (1D) are

$$\begin{aligned} \frac{\partial A}{\partial t} &= D_A \frac{\partial^2 u}{\partial x^2} + f(A,B) \\ \frac{\partial B}{\partial t} &= D_B \frac{\partial^2 v}{\partial x^2} + g(A,B) \end{aligned} \tag{18}$$

where $D_A$ and $D_B$ are the diffusion constants for the species $A$ and $B$. We assume that the above system has a stationary solution which is stable when there is no diffusion. Spatiotemporal stationarity needs to satisfy

$$\frac{\partial A}{\partial t} = 0, \quad \frac{\partial B}{\partial t} = 0, \quad \frac{\partial^2 A}{\partial x^2} = 0, \quad \frac{\partial^2 B}{\partial x^2} = 0 \tag{19}$$

Therefore

$$f(A,B) = 0, \quad g(A,B) = 0 \tag{20}$$

Depending on the functional form of $f(u,v)$ and $g(u,v)$, there might exist several stationary solutions, denoted by $(u_0, v_0)$.

##### 1.7.1 Stability

Linearize around the stationary solution and set $(A,B) = (A_0, B_0) + (\delta A(x,t), \delta B(x,t))$. Thus Eq.18 becomes

$$\frac{d}{dt}\begin{bmatrix} \delta A \\ \delta B \end{bmatrix} = \mathcal{S}\begin{bmatrix} \delta u \\ \delta v \end{bmatrix} + D\frac{\partial^2}{\partial x^2}\begin{bmatrix} \delta u \\ \delta v \end{bmatrix}$$

where the linear stability matrix and the diffusion matrix

$$\mathcal{S} = \begin{bmatrix} f_A & f_B \\ g_A & g_B \end{bmatrix}, \quad D = \begin{bmatrix} D_A & 0 \\ 0 & D_B \end{bmatrix}$$

Solutions of a linear PDE can be expressed as product of function of space and time, say, of the form

$$\begin{bmatrix} \delta A \\ \delta B \end{bmatrix} = e^{\lambda t + ikx}\mathbf{w_0}, \quad \mathbf{w_0} = \begin{bmatrix} \delta A_0 \\ \delta B_0 \end{bmatrix}$$

where $\delta A_0$ and $\delta B_0$ are constants. Thus 1.7.1 becomes

$$\lambda \mathbf{w}_0 = A\mathbf{w}_0 - Dk^2\mathbf{w}_0 \tag{21}$$

and the eigenvalues are determined by the solutions of the characteristic polynomial

$$Det(\mathcal{S} - Dk^2 - \lambda \mathbf{I}) = 0 \tag{22}$$

This gives eigenvalues $\lambda(k)$ as functions of wavenumber that are roots of the above equation, written explicitly as

$$\lambda^2 - \lambda\Big((f_u + g_v) - k^2(D_u + D_v)\Big) + h(k^2) = 0 \,,$$
$$h(k^2) = D_A D_B k^4 - (f_A D_B + g_B D_A)k^2 + f_A g_B - f_B g_A \tag{23}$$

The above equation is a quadratic equation in $k^2$, and since $D_A D_B > 0$, this is an equation of a convex parabola. We can calculate the wavenumber where the parabola is minimum (see next section). Thus the trace $\tau_{RD}$ and the determinant $\Delta_{RD}$ of the RD system is

$$\begin{aligned} \tau_{RD} &= (f_A + g_B) - k^2(D_A + D_B) \\ \Delta_{RD} &= h(k^2) = D_A D_B k^4 - (f_A D_B + g_B D_A)k^2 + f_A g_B - f_B g_A \end{aligned} \tag{24}$$

The steady state is linearly stable if both solutions of Eq.(23) satisfy $Re\lambda(k^2) < 0$. (Recollect that for Reaction systems (without diffusion), the condition for stability was $Re\lambda(k = 0) < 0$). Solving Eq.23, we get the eigenvalues $\lambda_{\pm}^{RD}(k^2)$ as functions of wave numbers, given by

$$2\lambda_{\pm}^{RD}(k^2) = (f_A + g_B) - k^2(D_A + D_B) \pm \left[\Big((f_A + g_B) - k^2(D_A + D_B)\Big)^2 - 4h(k^2)\right]^{1/2} \tag{25}$$

where $h(k^2)$ are the same functions as in Eq.23.

##### 1.7.2 Conditions for diffusive instability

For the steady state to become unstable with the diffusion present, we require
$Re\lambda^{RD}(k^2) > 0$ for some $k \neq 0$.
This can happen if either
1. the trace $\tau_{RD} > 0$, or if
2. the determinant $\Delta_{RD} = h(k^2) < 0$ for some $k \neq 0$.

We assumed $f_A + g_B < 0$ (trace of the stable Reaction system is negative) and $k^2(D_A + D_B) > 0$ for all $k \neq 0$. Hence for a RD system also, the trace, given by Eq(**??**) is always negative.

Thus to have instability with diffusion, we must have the determinant $\Delta_{RD} = h(k^2) = D_A D_B k^4 - (f_A D_B + g_B D_A)k^2 + f_A g_B - f_B g_A < 0$ for some $k \neq 0$. We know that the prefactor of the first term $D_A D_B > 0$ and the last term (determinant of the linear stability matrix of the Reaction system) $f_A g_B - f_B g_A > 0$. Thus, to ensure instability, i.e. for $\Delta < 0$, we need

$$f_A D_B + g_B D_A > 0 \tag{26}$$

We also know $f_A + g_B < 0$ (Trace of the linear stability matrix of the Reaction system is negative). So $D_A/D_B \neq 1$.

The inequality $(f_A D_B + g_B D_A) > 0$ is necessary, but not sufficient to have $Re(\lambda(k^2)) > 0$. For $\Delta_{RD} = h(k^2) < 0$ to satisfy for some $k \neq 0$, the minimum of $h(k^2)$ must be negative, i.e.,

$$\frac{\partial^2}{\partial k^2} h(k^2) = 2D_A D_B k^2 - D_B f_A - D_A g_B < 0 \tag{27}$$

Thus, the critical wave number is given by

$$k_c = \Big(\frac{D_A g_B + D_B f_A}{2D_A D_B}\Big)^{1/2} \tag{28}$$

This is the most unstable wave number and it corresponds to the fastest growing mode of patterns which occurs precisely at the bifurcation point where $min(h(k_c)) = 0$, $max(Re\lambda(k^2)) = 0$ and so

$$D_A g_B + D_B f_A = 4\big(D_A D_B (f_A g_B - f_B g_A)\big) \tag{29}$$

Inserting $k_c^2$ into $\Delta_{RD}$ gives the minimum value of the determinant.

$$\Delta_{RD}(k_c^2) = f_A g_B - f_B g_A - \frac{1}{4D_A D_B}\big(D_A g_B + D_B f_A\big)^2 \tag{30}$$

Now, to satisfy the condition for instability, we need the determinant $\Delta_c^{RD}(k_c^2)$ to be negative, which implies that the following equation should hold

$$D_A g_B + D_B f_A > 2\big(D_A D_B (f_A g_B - f_B g_A)\big)^{1/2} \tag{31}$$

In this regime, there exists a range of unstable wave numbers. The validity of the above equation also implies the validity of a less stringent condition, as follows

$$D_A g_B + D_B f_A > 0 \tag{32}$$

which is the same condition as expressed in Eq(26).

##### 1.7.3 Toggle Switch with 1D diffusion

The mathematical equations describing the spatiotemporal dynamics of toggle switches embedded in a one dimensional chain of cells are given by

$$\begin{aligned}
\frac{dA}{dt} &= g_A \mathcal{H}(B, B_0 A, n_{BA}, l_{BA}) - \gamma_A A + D_A \frac{\partial^2 A}{\partial x^2} \\
\frac{dB}{dt} &= g_B \mathcal{H}(A, A_0 B, n_{AB}, l_{AB}) - \gamma_B B + D_B \frac{\partial^2 B}{\partial x^2}
\end{aligned} \tag{33}$$

The trace, determinant and eigenvalues (Eq.s 24, 25) of the system can be determined by substituting $f_A$, $f_B$ and $g_A$, $g_B$ (as given in Eq.9). We notice that these expressions are functions of $k$ (variations of these quantities with $k$ is shown in Fig. 2 of the main text and Fig. S3 and S4 of SI).

##### 1.7.4 Toggle Switch with double self-activation with 1D diffusion

The mathematical equations describing the spatiotemporal dynamics of toggle switches with double self-activation embedded in a one dimensional chain of cells are given by

$$\begin{aligned}
\frac{dA}{dt} &= g_A\mathcal{H}(B, B_0A, n_{BA}, l_{BA})\mathcal{H}(B, A_0A, n_{AA}, l_{AA}) - \gamma_A A + D_A\frac{\partial^2 A}{\partial x^2} \\
\frac{dB}{dt} &= g_B\mathcal{H}(B, A_0B, n_{AB}, l_{AB})\mathcal{H}(A, B_0B, n_{BB}, l_{BB}) - \gamma_B B + D_B\frac{\partial^2 v}{\partial x^2}
\end{aligned} \tag{34}$$

The trace, determinant and eigenvalues (Eq.s 24, 25) of the system can be determined by substituting $f_A$, $f_B$ and $g_A$, $g_B$ (as given in Eq.12). We notice that these expressions are functions of $k$ (variations of these quantities with $k$ is shown in Fig. 3 of the main text and Fig. S5 of SI).

##### 1.7.5 Toggle Triad with 1D diffusion

The mathematical equations describing the spatiotemporal dynamics of toggle triads embedded in a one dimensional chain of cells are given by

$$\begin{aligned}
\frac{d\mathbf{A}}{dt} &= \mathcal{H}(\mathbf{B}, B_0A, n_{BA}, l_{BA})\mathcal{H}(\mathbf{C}, C_0A, n_{CA}, l_{CA}) - \gamma_A\mathbf{A} + D_A\frac{\partial^2 \mathbf{A}}{\partial x^2} \\
\frac{d\mathbf{B}}{dt} &= \mathcal{H}(\mathbf{C}, C_0B, n_{CB}, l_{CB})\mathcal{H}(\mathbf{A}, A_0B, n_{AB}, l_{AB}) - \gamma_B\mathbf{B} + D_B\frac{\partial^2 \mathbf{B}}{\partial x^2} \\
\frac{d\mathbf{C}}{dt} &= \mathcal{H}(\mathbf{A}, A_0C, n_{AC}, l_{AC})\mathcal{H}(\mathbf{B}, B_0C, n_{BC}, l_{BC}) - \gamma_C\mathbf{C} + D_C\frac{\partial^2 \mathbf{C}}{\partial x^2}
\end{aligned} \tag{35}$$

The trace, determinant and eigenvalues **??** of the system can be determined by substituting $f_A$, $f_B$, $f_C$ and $g_A$, $g_B$, $g_C$ (as given in Eq.**??**). We notice that these expressions are functions of $k$ (variations of these quantities with $k$ is shown in Fig. **??** of the main text).

#### 1.8 Reaction-Diffusion Systems in 2D

The mathematical equations representing a 2-component (u,v) RD system in 2-dimensional space are given by

$$\begin{aligned}
\frac{\partial u}{\partial t} &= D_u^x\frac{\partial^2 u}{\partial x^2} + D_u^y\frac{\partial^2 u}{\partial y^2} + f(u, v) \\
\frac{\partial v}{\partial t} &= D_v^x\frac{\partial^2 v}{\partial x^2} + D_v^y\frac{\partial^2 v}{\partial y^2} + g(u, v)
\end{aligned} \tag{36}$$

Thus the trace $\tau_{RD}$, the determinant $\Delta_{RD}$ and the eigenvalues of the RD system in two dimension is given by

$$
\begin{aligned}
\tau_{RD} &= (f_u + g_v) - k^2(D_u^x + D_u^y + D_v^x + D_v^y) \\
\Delta_{RD} &= h(k^2) = D_u^x D_u^y D_v^x D_v^y k^4 - (f_u(D_v^x + D_v^y) + g_v(D_u^x + D_u^y))k^2 + f_u g_v - f_v g_u \\
2\lambda_{\pm}^{RD}(k^2) &= (f_u + g_v) - k^2(D_u^x + D_u^y + D_v^x + D_v^y) \\
&\pm\left[\left((f_u + g_v) - k^2(D_u^x + D_u^y + D_v^x + D_v^y)\right)^2 - 4h(k^2)\right]^{1/2}
\end{aligned} \tag{37}
$$

##### 1.8.1 Toggle Switch with 2D diffusion

The mathematical equations which can describe toggle switch with molecules diffusing in two dimensional space

$$
\begin{aligned}
\frac{dA}{dt} &= = g_A \mathcal{H}(B, B_0 A, n_{BA}, l_{BA}) - \gamma_A A + D_A^x \frac{\partial^2 A}{\partial x^2} + D_A^y \frac{\partial^2 A}{\partial y^2} \\
\frac{dB}{dt} &= = g_B \mathcal{H}(A, A_0 B, n_{AB}, l_{AB}) - \gamma_B B + D_B^x \frac{\partial^2 B}{\partial x^2} + D_B^y \frac{\partial^2 B}{\partial y^2}
\end{aligned} \tag{38}
$$

The trace, determinant and eigenvalues, given by Eq. 37 of the system can be determined by substituting $f_A$, $f_B$ and $g_A$, $g_B$ (as given in Eq.9). We notice that these expressions are functions of $k$ (variations of these quatities with $k$ is shown in Fig. 2 of the main text and Fig. S3 and S4 of SI).

##### 1.8.2 Toggle Switch with double self-activation with 2D diffusion

The mathematical equations which can describe toggle switch with double self-activation loops with molecules diffusing in two dimensional space

$$
\begin{aligned}
\frac{du}{dt} &= g_A \mathcal{H}(v, B_0 A, n_{BA}, l_{BA}) \mathcal{H}(v, A_0 A, n_{AA}, l_{AA}) - \gamma_A u + D_u^x \frac{\partial^2 u}{\partial x^2} + D_u^y \frac{\partial^2 u}{\partial y^2} \\
\frac{dv}{dt} &= g_B \mathcal{H}(u, A_0 B, n_{AB}, l_{AB}) \mathcal{H}(u, B_0 B, n_{BB}, l_{BB}) - \gamma_B v + D_v \frac{\partial^2 v}{\partial x^2} + D_v^y \frac{\partial^2 v}{\partial y^2}
\end{aligned} \tag{39}
$$

The trace, determinant and eigenvalues (Eq. 37) of the system can be determined by substituting $f_A$, $f_B$ and $g_A$, $g_B$ (as given in Eq.12). We notice that these expressions are functions of $k$ (variations of these quantities with $k$ is shown in Fig. 3 of the main text and Fig. S5 of SI).

##### 1.8.3 Toggle Triad with 2D diffusion

The mathematical equations which can describe toggle triad with molecules diffusing in two dimensional space

$$
\begin{aligned}
\frac{d\mathbf{A}}{dt} &= \mathcal{H}(\mathbf{B}, B_0A, n_{BA}, l_{BA})\mathcal{H}(\mathbf{C}, C_0A, n_{CA}, l_{CA}) - \gamma_A\mathbf{A} + D_A^x\frac{\partial^2\mathbf{A}}{\partial x^2} + D_A^y\frac{\partial^2\mathbf{A}}{\partial y^2} \\
\frac{d\mathbf{B}}{dt} &= \mathcal{H}(\mathbf{C}, C_0B, n_{CB}, l_{CB})\mathcal{H}(\mathbf{A}, A_0B, n_{AB}, l_{AB}) - \gamma_B\mathbf{B} + D_B^x\frac{\partial^2\mathbf{B}}{\partial x^2} + D_B^y\frac{\partial^2 B}{\partial y^2} \\
\frac{d\mathbf{C}}{dt} &= \mathcal{H}(\mathbf{A}, A_0C, n_{AC}, l_{AC})\mathcal{H}(\mathbf{B}, B_0C, n_{BC}, l_{BC}) - \gamma_C\mathbf{C} + D_C^x\frac{\partial^2\mathbf{C}}{\partial x^2} + D_C^y\frac{\partial^2\mathbf{C}}{\partial y^2} \quad (40)
\end{aligned}
$$

The trace, determinant and eigenvalues of the system can be determined by substituting $f_A$, $f_B$, $f_C$ and $g_A$, $g_B$, $g_C$ (as given in Equations 14, 15 and 16). We notice that the final expressions will be functions of the wavenumber $k$ (variations of these quantities with $k$ is shown in Figures S8, S9, and S10 of SI).

#### 2 Supplementary Figures

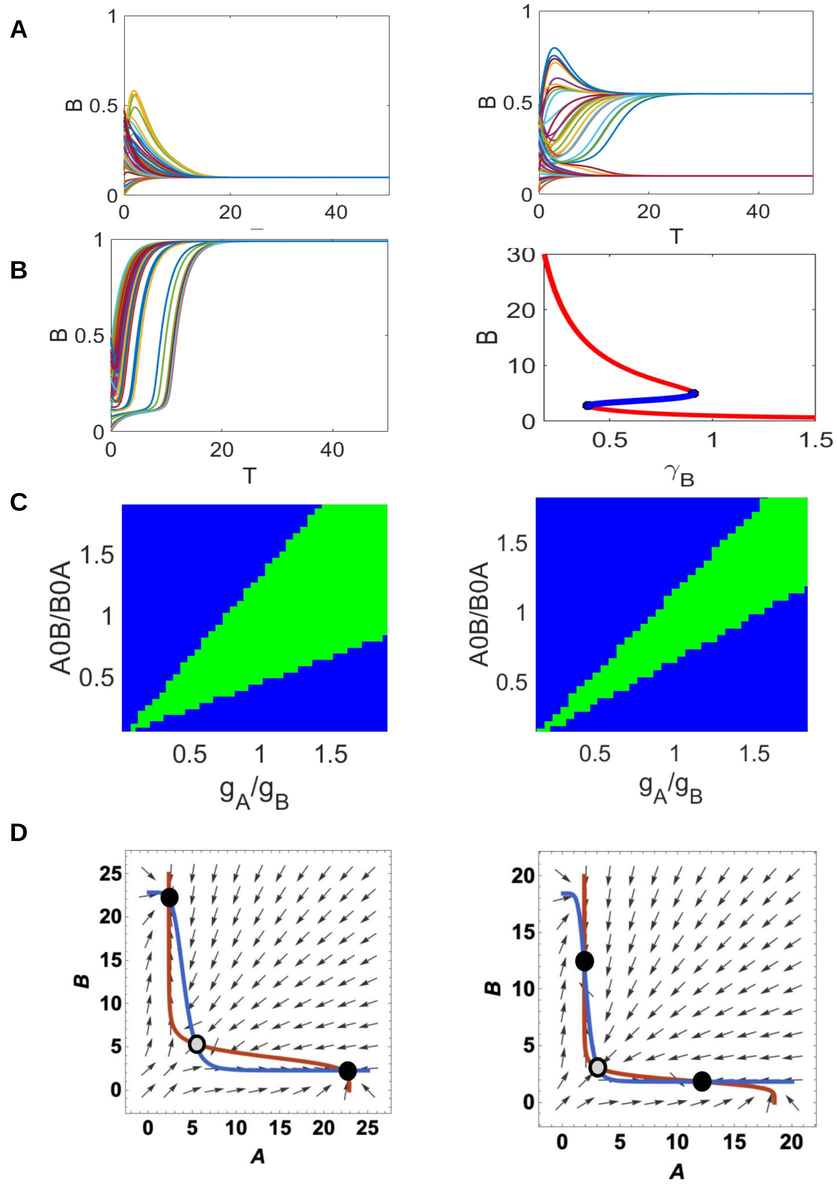


Figure S1: A and B) Dynamics and bifurcation diagram for the gene B for the parameter set 1 for TS shown in Figure 1. C) Phase diagram for two other parameter sets (2 and 3) depicting the behaviour of toggle switch. D) Nullclines from the bistable regions of the phase diagrams for parameter set 2 and 3.

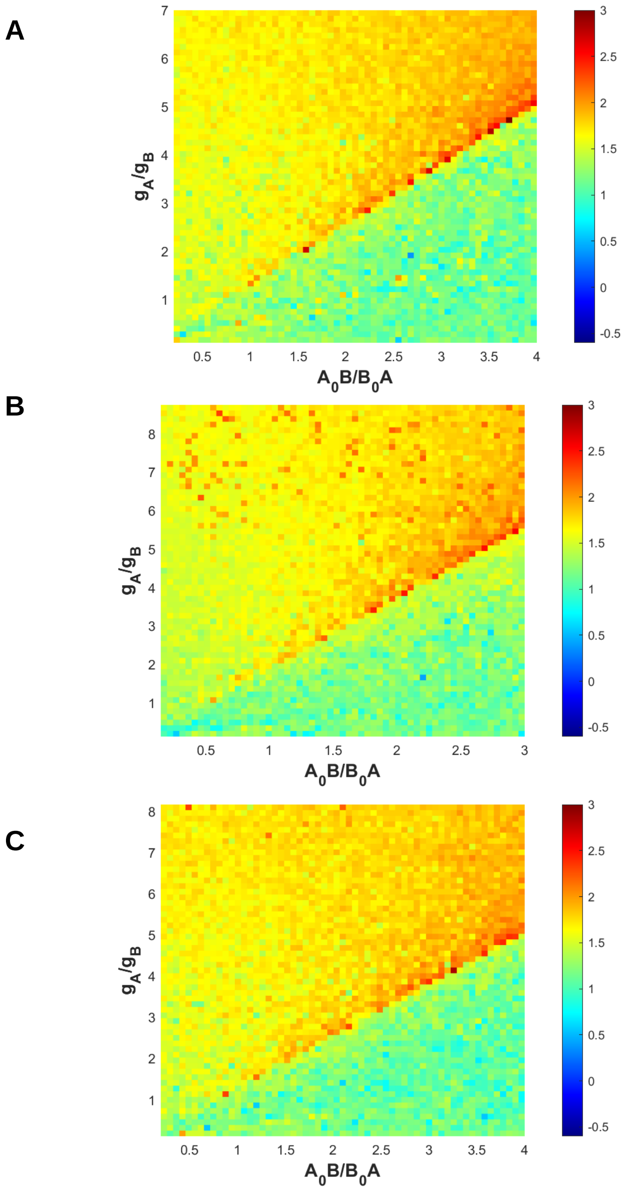


Figure S2: Phase Diagrams for the time (shown in logarithmic scale) taken by gene A to reach the steady state for different parameter regimes for parameter sets : A) 1 B) 2 and C) 3 for Toggle Switch.

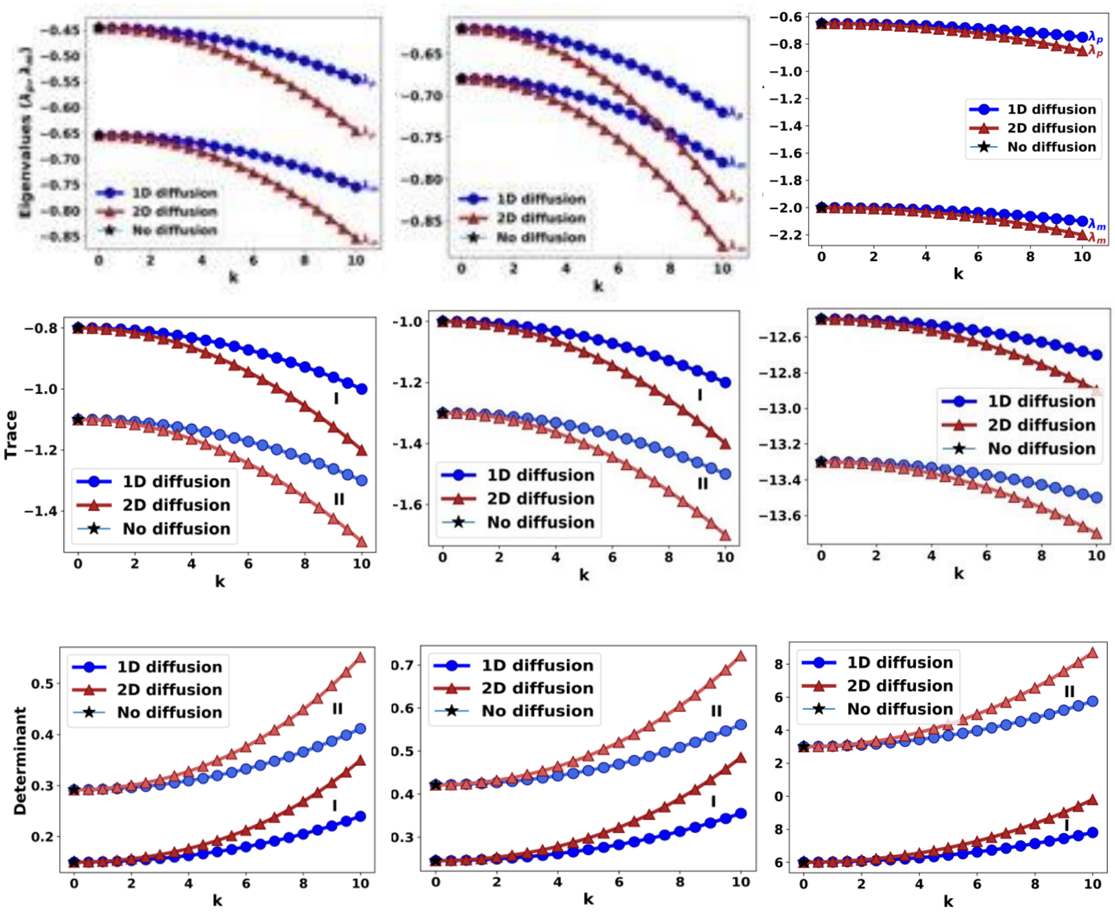


Figure S3: A) Eigenvalues corresponding to three different regimes - Monostable I, Bistable II and Monostable III of Toggle Switch corresponding to parameter set 2, B) Traces for three different regimes of parameter set 1 and 2, C) Determinants for three different regimes of parameter set 1 and 2

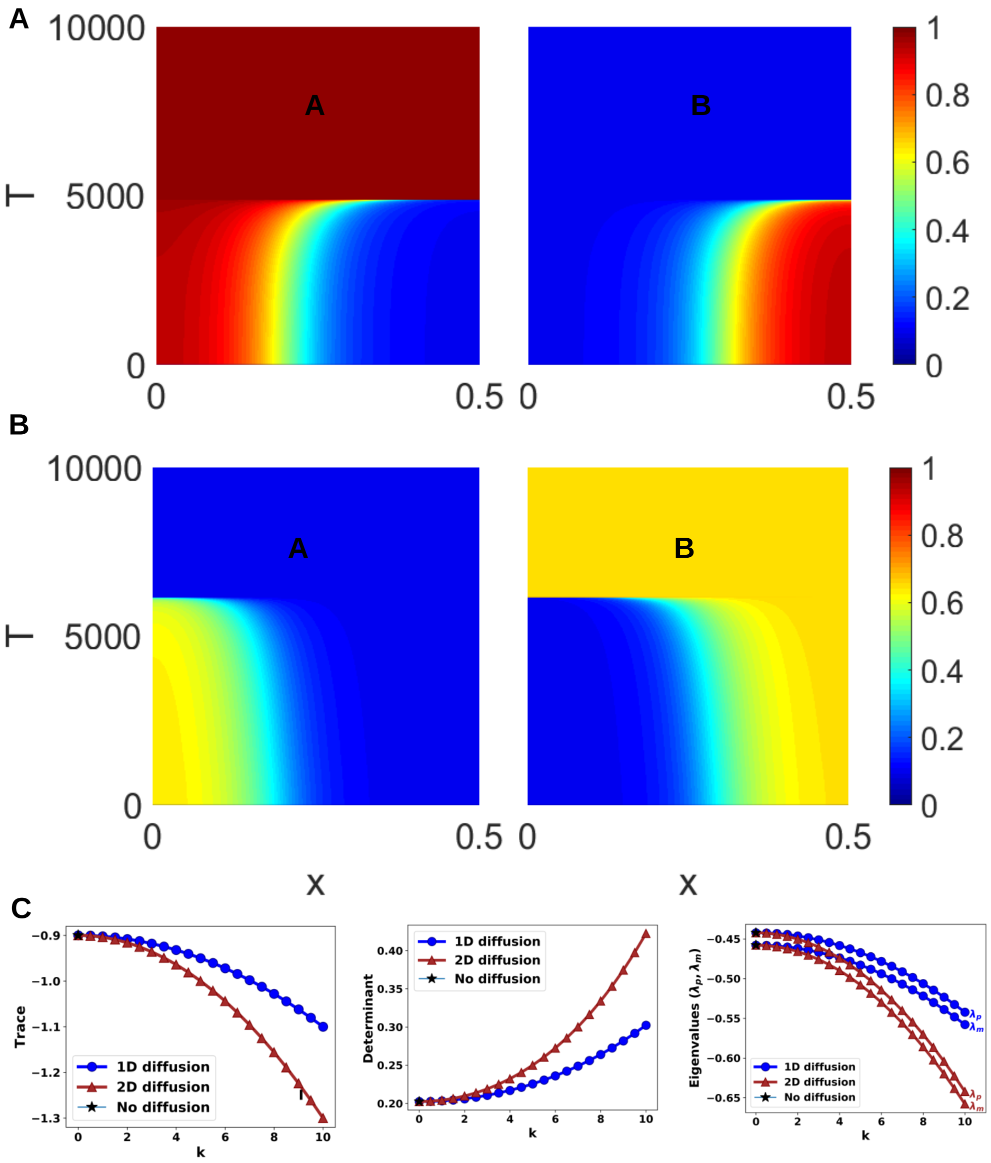


Figure S4: A) Heatmaps of expression levels of gene A and gene B as a function of space and time for parameter set 2 of Toggle Switch with 1D diffusion B) Heatmaps of expression levels of gene A and gene B as a function of space and time for parameter set 3 C) Trace, Determinant and Eigenvalues for parameter set 3

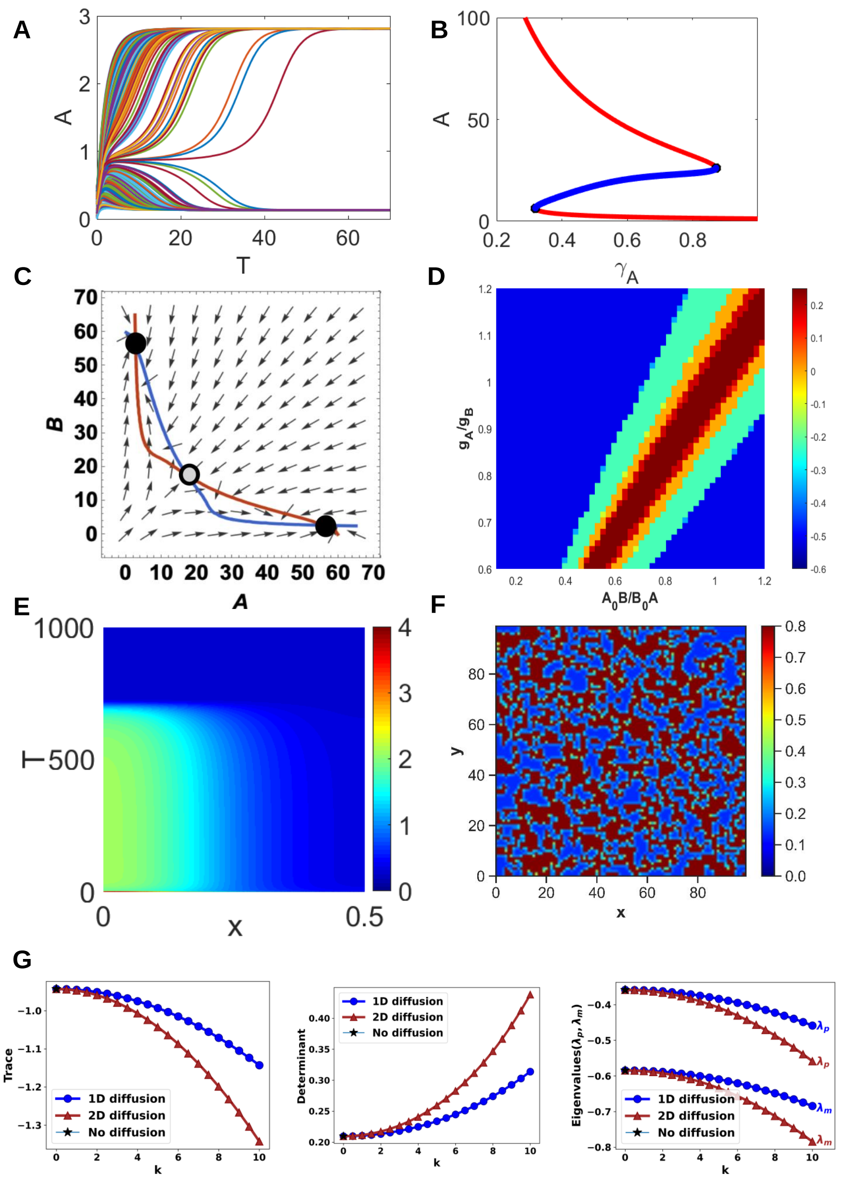


Figure S5: Bistable regime in TSSA: A) Time evolution plot of protein A for multiple initial conditions in the bistable parameter regime. B) Bifurcation diagram of A as a function of degradation rate of A. C) Nullcline plot of A and B in the bistable regime. D) Phase Diagram of log of the time it takes to reach the steady state for different parameter regimes. E) Heatmap of levels of A as a function of space and time for the 1D diffusion case F) ) Heatmap of levels of A as a function of space for the 2D diffusion case G) Trace, Determinant and Eigenvalues for the bistable regime of the parameter set A

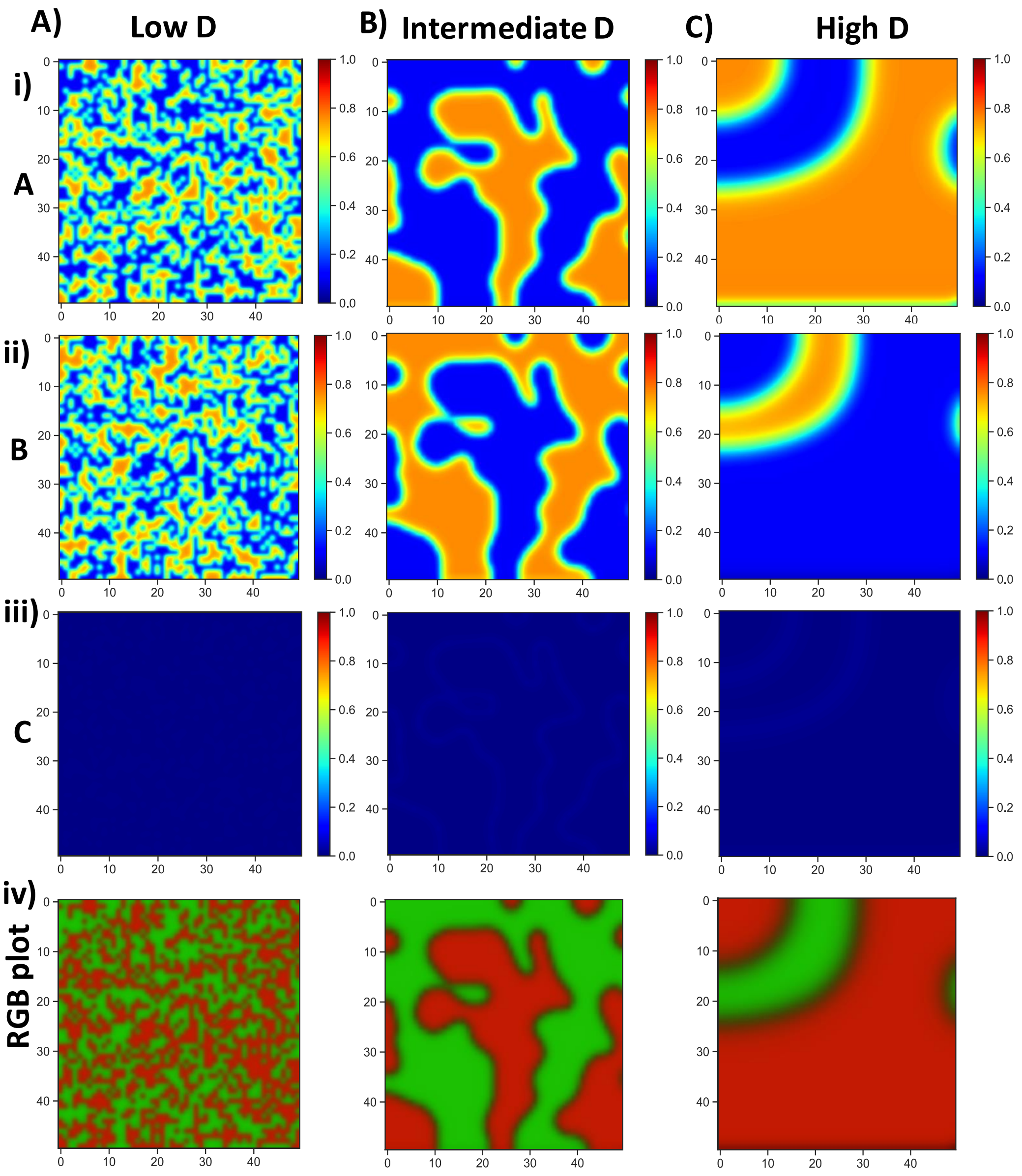


Figure S6: Dynamics of Toggle Triad in the Bistable parameter regime for different diffusion constants. A) Heat map of levels of i) A ii) B iii) C as a function of space in the bistable parameter regime when the diffusion is low (D=0.0008). iv) RGB plot of the 3 species combined. Here, red corresponds to A high, green corresponds to B high, and blue corresponds to C high. B) Same as A but for intermediate diffusion levels (D=0.008) C) Same as A but for high diffusion levels (D=0.08)

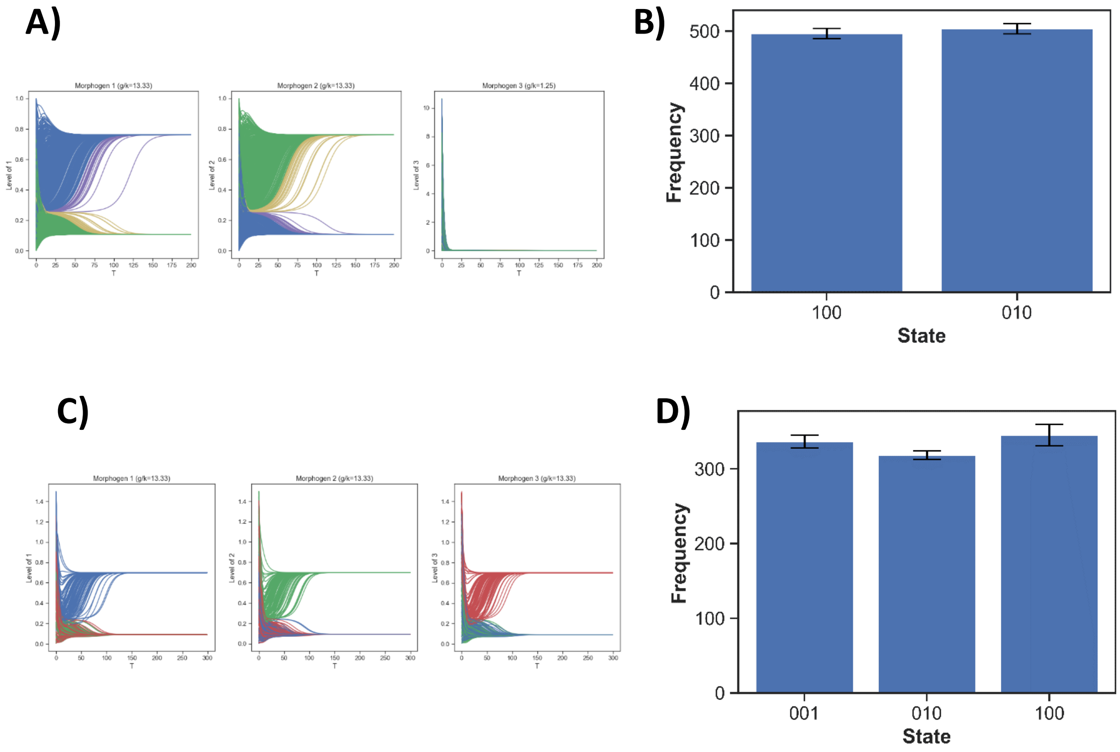


Figure S7: A) Time evolution plot of molecules of gene A for multiple initial conditions in the bistable parameter regime of toggle triad. B) Frequency distribution of different states at steady state. Error bars are calculated after three independent runs. C) Time evolution plot of protein A for multiple initial conditions in the tristable parameter regime. D) Frequency distribution of different states at steady state. Error bars are calculated after three independent runs.

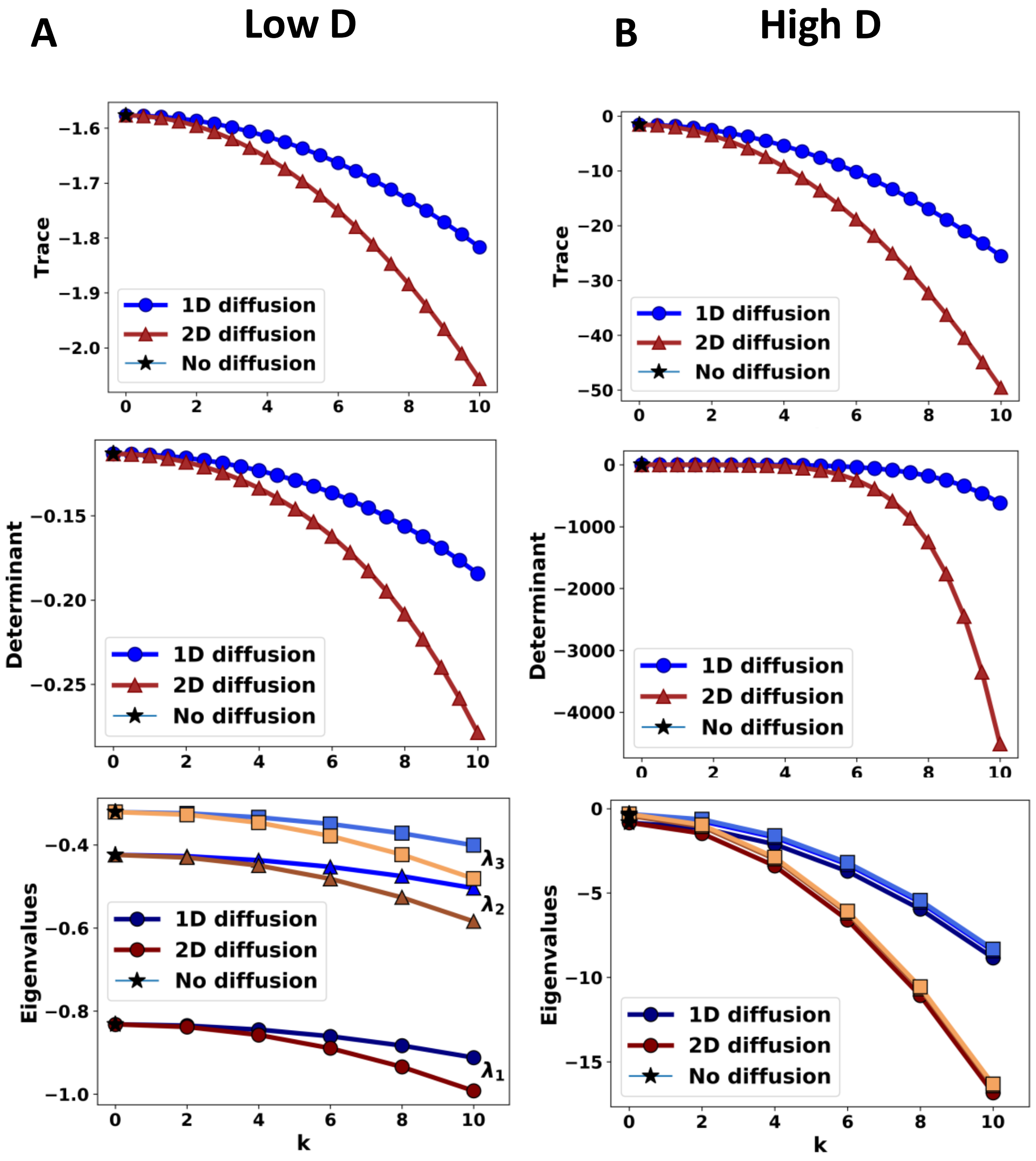


Figure S8: Stability analysis for Toggle Triad (monostable parameter regime). Column A shows Trace, Determinant and Eigenvalues for low diffusion coefficient for TT without diffusion (black) and with 1D (blue) and 2D diffusion (brown). Column B shows Trace, Determinant and Eigenvalues for high diffusion coefficient for TT without diffusion (black) and with 1D (blue) and 2D diffusion (brown). With the increase in $k$, the trace, determinant and eigenvalues become more and more negative. With high diffusion coefficient, the quantities reach much higher negative values at a faster rate (see the ranges in the y-axis)

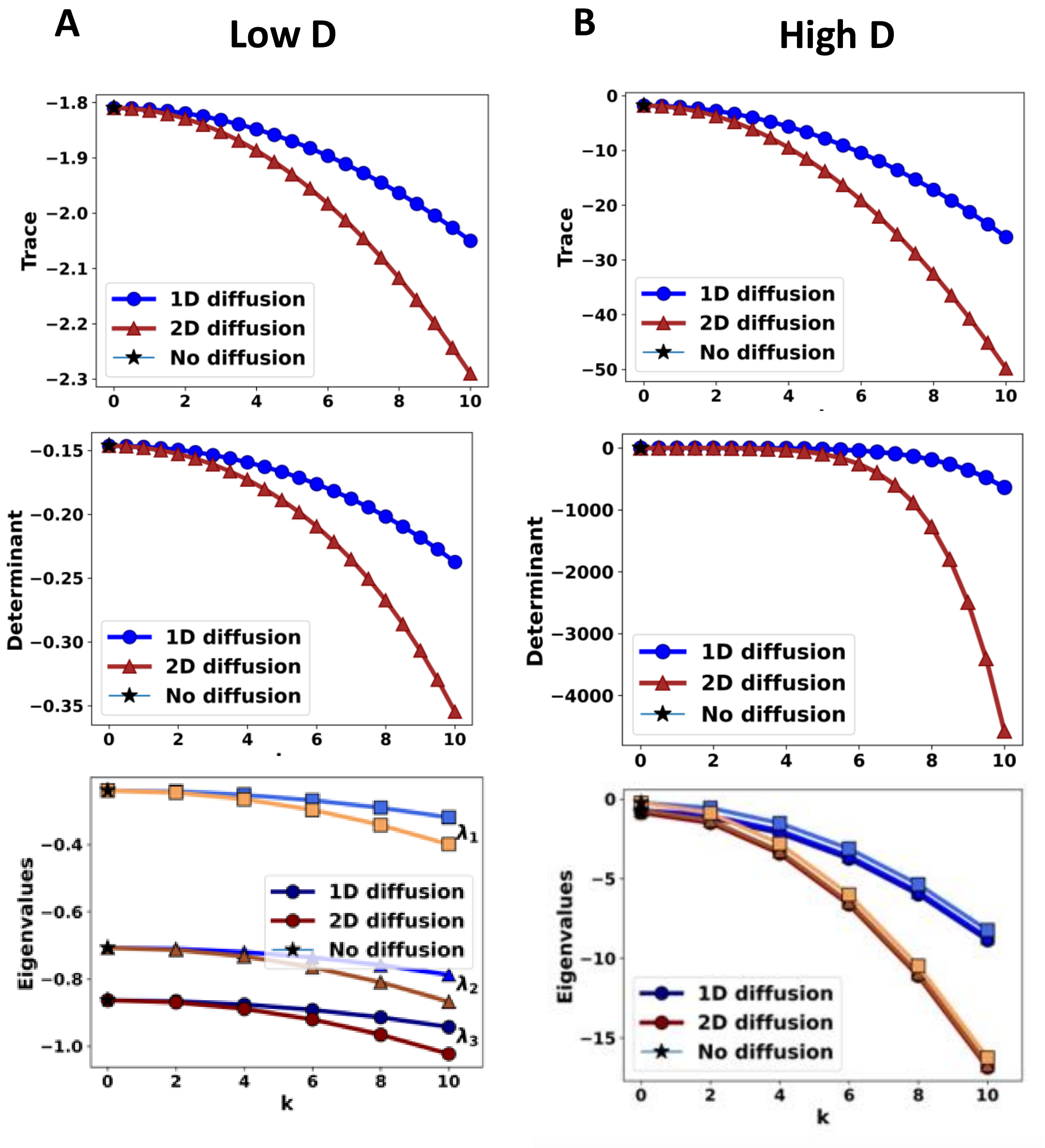


Figure S9: Stability analysis for Toggle Triad (bistable parameter regime). Column A shows Trace, Determinant and Eigenvalues for low diffusion coefficient for TT without diffusion (black) and with 1D (blue) and 2D diffusion (brown). Column B shows Trace, Determinant and Eigenvalues for high diffusion coefficient for TT without diffusion (black) and with 1D (blue) and 2D diffusion (brown). With the increase in $k$, the trace, determinant and eigenvalues become more and more negative. With high diffusion coefficient, the quantities reach much higher negative values at a faster rate (see the ranges in the y-axis). Exactly same result is obtained when these quantities are evaluated at the second equilibrium point, since the coordinates of the equilibrium points are in symmetric pairs. We notice that the ranges of y-axes in the plots for trace, determinants and eigenvalues are very similar to one another in the mono-, bi- and tristable regimes. This is because at high diffusion coefficient, the system stabilizes/homogenizes very fast independent of the other parameter values.

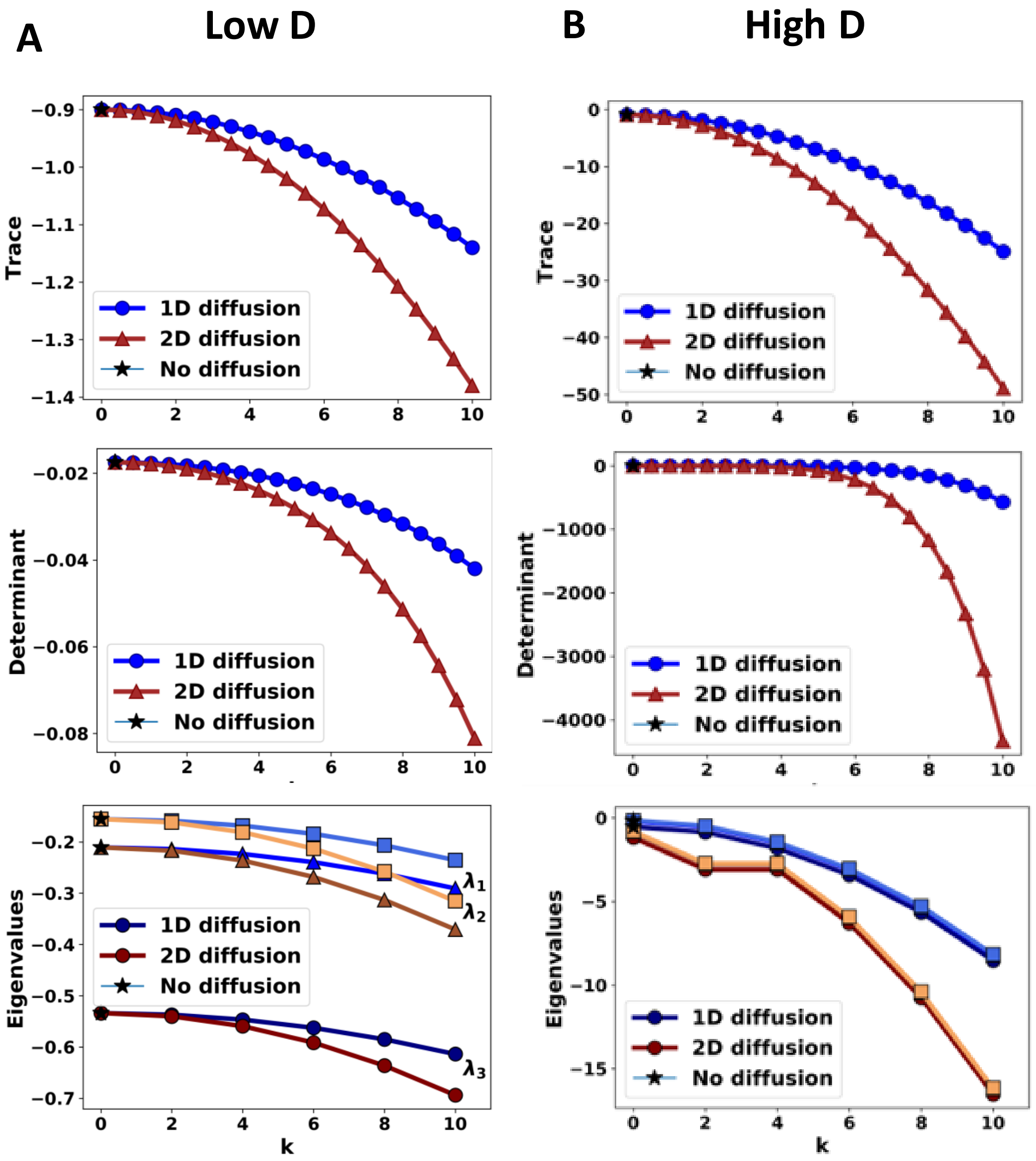


Figure S10: Stability analysis for Toggle Triad (tristable parameter regime). Column A shows Trace, Determinant and Eigenvalues for low diffusion coefficient for TT without diffusion (black) and with 1D (blue) and 2D diffusion (brown). Column B shows Trace, Determinant and Eigenvalues for high diffusion coefficient for TT without diffusion (black) and with 1D (blue) and 2D diffusion (brown). With the increase in $k$, the trace, determinant and eigenvalues become more and more negative. With high diffusion coefficient, the quantities reach much higher negative values at a faster rate (see the ranges in the y-axis). Exactly same results are obtained when these quantities are evaluated at the second and third equilibrium points, since the coordinates of the equilibrium points are in symmetric triads

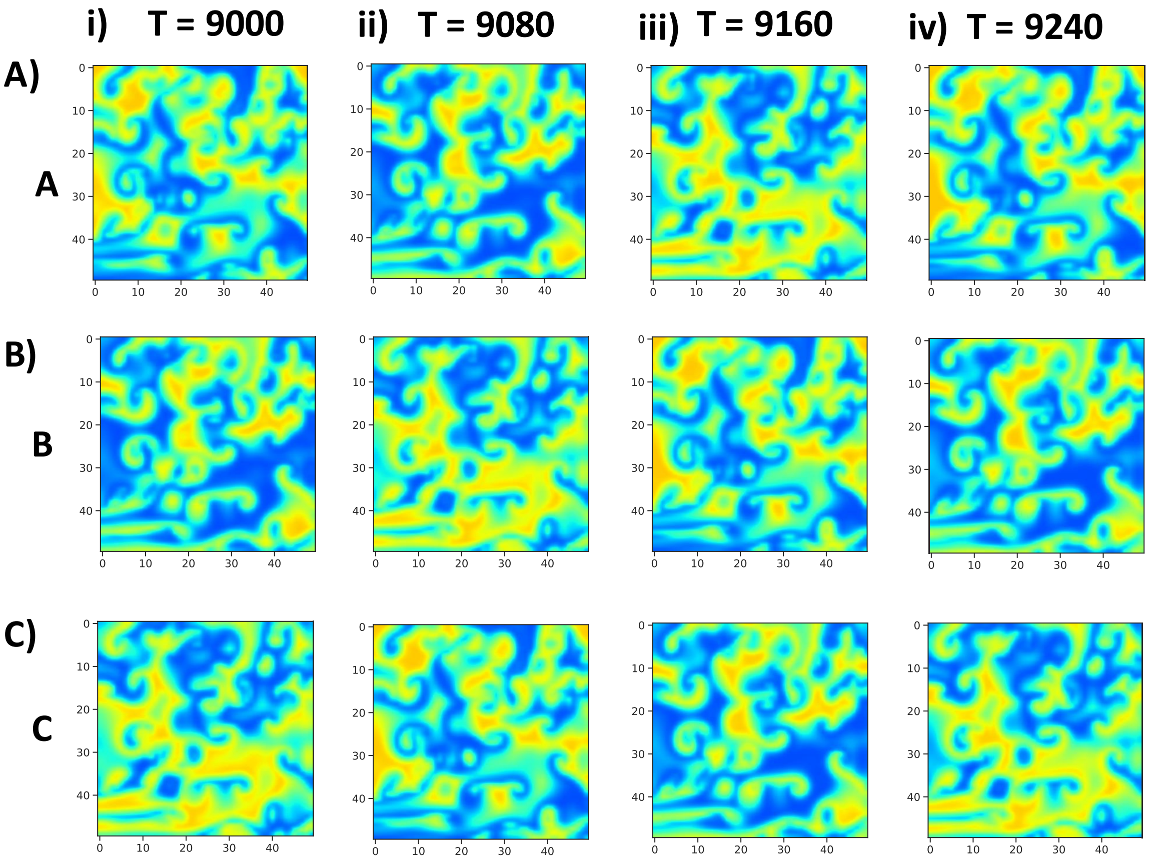


Figure S11: Repressilator: Heatmaps of levels of A) gene A, B) gene B and C) gene C as a function of space at four different time-points ( i) T = 9000, ii) T = 9080 iii) T = 9160 iv) T = 9240)

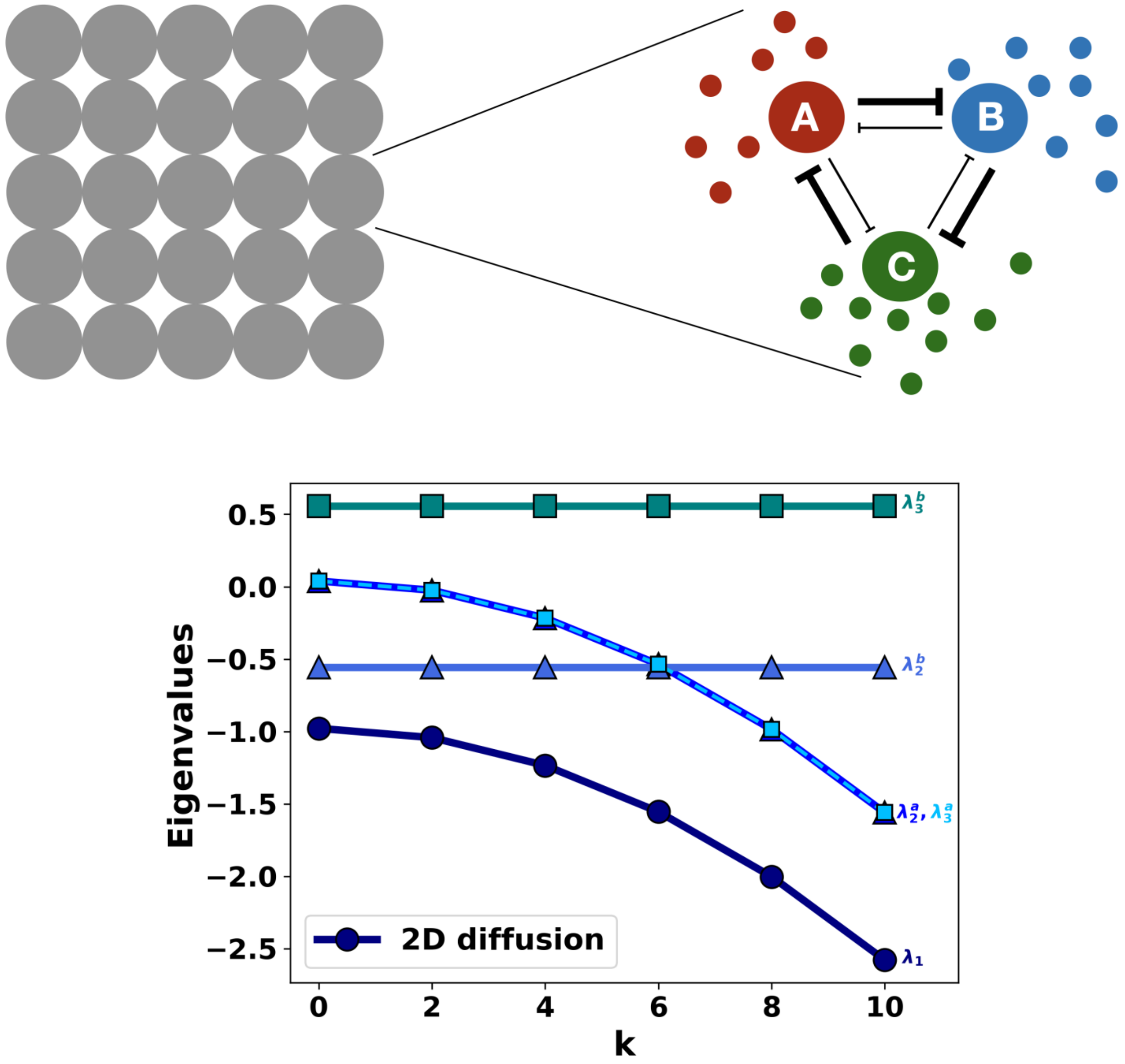


Figure S12: Analytical analysis of repressilator: The system has three eigenvalues, viz. $\lambda_1$ (shown in circles), $\lambda_2 = \lambda_2^a + i\lambda_2^b$ (shown in triangles) and $\lambda_3 = \lambda_3^a + i\lambda_3^b$ (shown in squares). $\lambda_1$ is negative for all values of $k$ ($\lambda_1 < 0\ \forall k$), $\lambda_2$ is complex, with positive real part $\lambda_2^a > 0$ for $k = 0$. For $k > 0$, it is negative i.e. $\lambda_2^a < 0$ while the imaginary part $\lambda_2^b < 0\ \forall k$ and has the same magnitude for all $k$. $\lambda_2$ and $\lambda_3$ are complex conjugate of each other. Thus the straight lines in the plot are the imaginary parts which does not vary with $k$.

### 3 Supplementary videos

1. Animation of Toggle switch coupled with diffusion in 2D space Video of the simulation represented in Fig 2 C. In this simulation, we coupled toggle switch with diffusion in a 2D lattice. We observe patches of A high or B high which grow over time. Depending on the initial conditions the system either goes to homogeneous A high state or B high state.

2. Animation of Toggle switch plus Self-Activation coupled with diffusion in 2D space

   The video of the simulation is represented in Fig 2 C. We coupled the toggle switch with diffusion in a 2D lattice in this simulation. We observe patches of A high or B high which grow over time. Depending on the initial conditions, the system either goes to a homogeneous A high state or B high state.

3. Animation of Toggle Triad coupled with diffusion in 2D space

   Video of the simulation represented in Fig 5 C for the Toggle Triad in the tristable regime. The system evolves from random initial conditions and patches of "Abc" or "aBc" or "abC". Depending on the initial conditions, the system either goes to a a homogeneous A high state, B high state, or C high state.

4. Animation of Repressilator coupled with diffusion in 2D space Video of the simulation represented in Fig 6 (Toggle switch plus self activation). In this simulation, we coupled an oscillatory motif- repressilator with diffusion in a 2D lattice. After a certain time point (around 1 min here), the pattern repeats itself. We observe the local spiral patterns that repeat themselves.
